# Dynamics and structural changes of calmodulin upon interaction with its potent antagonist calmidazolium

**DOI:** 10.1101/2022.01.19.474921

**Authors:** Corentin Léger, Irène Pitard, Mirko Sadi, Nicolas Carvalho, Sébastien Brier, Ariel Mechaly, Dorothée Raoux-Barbot, Maryline Davi, Sylviane Hoos, Patrick Weber, Patrice Vachette, Dominique Durand, Ahmed Haouz, J. Iñaki Guijarro, Daniel Ladant, Alexandre Chenal

## Abstract

Calmodulin (CaM) is a eukaryotic multifunctional, calcium-modulated protein that regulates the activity of numerous effector proteins involved in a variety of physiological processes. Calmidazolium (CDZ) is a potent small molecule antagonist of CaM and one the most widely used inhibitors of CaM in cell biology. Here, we report the structural characterization of CaM:CDZ complexes using combined SAXS, X-ray crystallography, HDX-MS and NMR approaches. Our results provide molecular insights into the CDZ-induced dynamics and structural changes of CaM leading to its inhibition. CDZ-binding induces an open-to-closed conformational change of CaM and results in a strong stabilization of its structural elements associated with a reduction of protein dynamics over a large time range. These CDZ-triggered CaM changes mimic those induced by CaM-binding peptides derived from protein targets, despite their distant chemical nature. CaM residues in close contact with CDZ and involved in the stabilization of the CaM:CDZ complex have been identified. These results open the way to rationally design new CaM-selective drugs.

**Figure and text for the Table of Contents (ToC):** 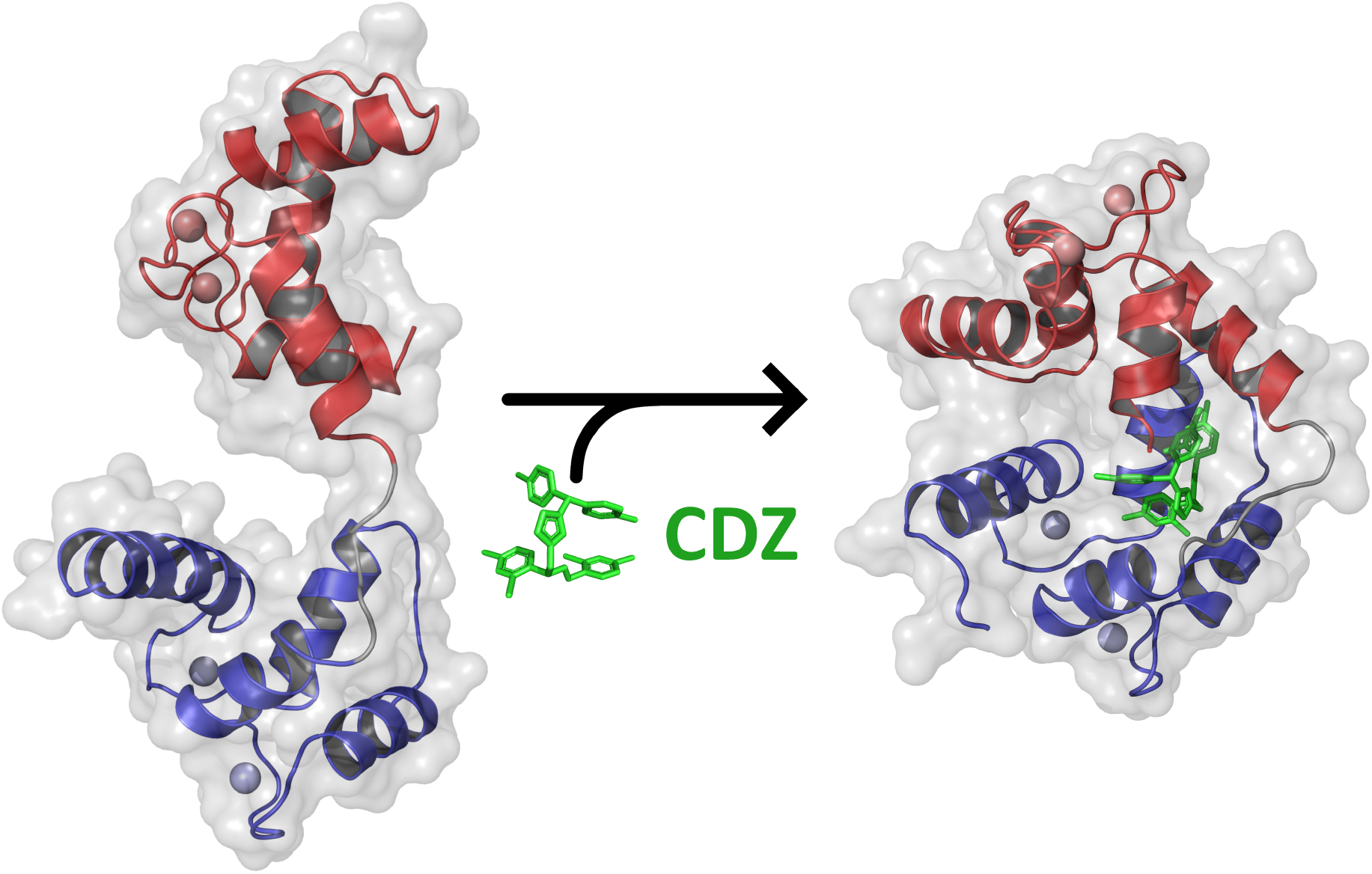

Calmidazolium is a potent and widely used inhibitor of calmodulin, a major mediator of calcium-signaling in eukaryotic cells. Structural characterization of calmidazolium-binding to calmodulin reveals that it triggers open-to-closed conformational changes similar to those induced by calmodulin-binding peptides derived from enzyme targets. These results open the way to rationally design new and more selective inhibitors of calmodulin.

## 1. Introduction

Calmodulin (CaM) is an essential 148 amino acid-long calcium binding protein that is ubiquitously found in all eukaryotes and one of the most conserved proteins known to date, playing a central role in cell physiology as a key sensor of intracellular calcium signaling ^1–3^. It regulates a wide variety of biochemical processes by interacting in a calcium-dependent manner with numerous effector proteins to modulate their enzymatic activities and/or their structural properties. Calmodulin has four calcium binding sites, called EF-hand motifs, made of about 30 amino acids that adopt a helix-loop-helix fold ^4^. Calcium binds with high affinity to the loop flanked by the two helical segments. CaM has two pairs of EF-hands (N-lobe and C-lobe) connected by a flexible linker. Calcium binding to EF-hands triggers large conformational changes that result in the exposure of hydrophobic patches and promote interaction with target proteins ^5,6^. Structural studies have highlighted the remarkable plasticity of CaM upon association with its effectors. In many instances, CaM binds its target proteins *via* short, 20–30 residues long, CaM-binding sites (CBS) with a positively charged, amphipathic alpha-helical character. Calcium-bound CaM (holo-CaM) generally binds by wrapping its two lobes around the CBS helix, adopting widely diverse configurations that allow it to match the great variability of the primary amino-acid sequences of CBS ^7,1,8,9^. For many effectors, the CBS sequence is located close to an auto-inhibitory domain that maintains the target enzyme in an inhibited state until holo-CaM binding to the CBS relieves the inhibition ^10,9^. Alternatively, holo-CaM can interact with targets in a noncanonical fashion in which the N- and C-lobes can simultaneously and independently associate to either distinct sites on the same protein or to the same site on two distinct polypeptide chains, triggering oligomerization ^10–12^. Besides, CaM can also associate with various proteins regardless of the presence of calcium, frequently *via* an ‘IQ motif’ (IQXXXRGXXXR) that interacts with both calcium-bound and calcium-free CaM (Apo-CaM) ^13,10^.

CaM mediates many physiological processes such as inflammation, metabolism, proliferation, apoptosis, smooth muscle contraction, intracellular movement, short-term and long-term memory, immune responses, etc ^14,15,2,9,12^. Given its central role in cell physiology, drugs able to antagonize CaM activity are in dire need for fundamental studies of calcium/CaM signaling as well as for potential therapeutic applications. A variety of molecules with diverse chemical structures were characterized in the 80’s following the initial observations that the phenothiazine family of antipsychotic drugs could inhibit the actions of holo-CaM ^16–21^. Many pharmacologically active drugs (antidepressants, anticholinergics, smooth muscle relaxants, local anesthetics, antiparasitic) were also shown to bind holo-CaM. It was suggested that their therapeutic activity might be, in part, related to their ability to interfere with the calcium/CaM signaling axis, which was therefore seen as a potentially relevant pharmacological target ^21,22^. At a fundamental level, these drugs have also been widely used to probe the role of CaM in cell biology. A major issue with these CaM antagonists is their lack of selectivity as most of them also target multiple additional effectors, making difficult to ascribe specific effects to their calcium/CaM signaling pathway inhibition ^23,24^. Although these CaM antagonists display a variety of chemical structures, they generally share overall hydrophobicity and a net positive charge. Their CaM-binding affinities as well as their stoichiometries are also variable. Structures of several of these antagonists in complex with CaM have revealed the diversity in their protein binding mode, akin to the variability of CaM-CBS complexes ^25–30^.

Here we provide a detailed characterization of the structural and dynamic changes of holo-CaM induced by the binding of calmidazolium (CDZ), one of the most potent and widely used CaM antagonists ^19,31–33,22,34^. We demonstrate both in crystal and in-solution that the binding of a single CDZ, primarily to the N-lobe of holo-CaM, is enough to induce an open-to-closed conformational reorganization associated with drastic changes of CaM dynamics. This is in marked contrast to prior studies that speculated that multiple CDZ could bind a single CaM molecule. We further delineate the structural rearrangements and changes in internal dynamics triggered by CDZ binding to CaM, as well as the dynamics of the association. These structural data will be instrumental to design CDZ analogs with improved selectivity for CaM.

## 2. Results

### 2.1. CDZ binding to holo-CaM monitored by SRCD and ITC

We first investigated the effect of CDZ on the secondary structure content of CaM using circular dichroism in the far-UV range to determine whether CDZ binding impacts CaM folding (Figure S1). The addition of CDZ does not induce major changes of the far-UV CD spectrum of calcium-loaded calmodulin (holo-CaM), suggesting that CDZ binding does not significantly alter the secondary structure content of holo-CaM. We then investigated the thermodynamics of CDZ binding to holo-CaM by isothermal titration calorimetry (ITC). The data indicated an apparent equilibrium dissociation KD constant of 3 ± 2 μM with a stoichiometry of 1.2 ± 0.5 (Figure S2), in agreement with Dagher et al. ^33^. An integrative structural biology approach combining SAXS, X-ray crystallography (XR), HDX-MS and NMR was then used to provide molecular insights into the CDZ-induced dynamics and conformational changes of holo-CaM, leading to its inhibition. ^31,19,32,33,22,34,35^

### 2.2. Analysis of CDZ-induced holo-CaM conformational changes by SEC-SAXS

We analyzed the effects of CDZ binding on the molecular shape of holo-CaM by Size exclusion chromatography coupled to small-angle X-ray scattering (SEC-SAXS) measurements. ^36^ The experiments were performed in the presence of 5% DMSO to prevent CDZ aggregation (see Supplementary info for details). The SEC-SAXS patterns recorded for holo-CaM in the absence and in the presence of 5% DMSO were essentially identical, suggesting that 5% DMSO has no detectable effects on holo-CaM (Figure S3).

The SEC-SAXS patterns of holo-CaM in the absence and in the presence of CDZ are shown in Figure 1A and the derived structural parameters are reported in Table S1. The SAXS patterns exhibit dramatic differences that are further highlighted by the pair-distance distribution function, P(r), and the dimensionless Kratky analysis (Figures 1B and 1C). The distance distribution functions show that holo-CaM adopts a bi-lobed conformation (SASDNX3) and that CDZ binding induces a compaction of the protein, leading to a globular shape (SASDNY3). The CDZ-induced conformational change of holo-CaM is characterized by a dramatic reduction of the radius of gyration, R_g_, from 22.4 to 16.8 Å and of the maximal interatomic distances, D_max_, from 72 Å to 52 Å (Figure 1B and Table S1). The dimensionless Kratky plots indicate a significant reduction of the structural flexibility of holo-CaM upon CDZ binding (Figure 1C), with the holo-CaM:CDZ complex exhibiting the archetypical profile of a folded, compact and isometric protein. *Ab initio* models of CaM global conformations were calculated using DENSS^37^, yielding a bi-lobed extended shape for holo-CaM and a globular shape for the holo-CaM:CDZ complex (Figures 1D and 1E), in agreement with the pair-distance distribution functions and the dimensionless Kratky plots (Figures 1B and 1C). Taken together, the SAXS results indicate that CDZ-binding induces an open-to-closed conformational change and a reduction of holo-CaM flexibility, illustrating the conformational plasticity of holo-CaM.

**Figure 1.**
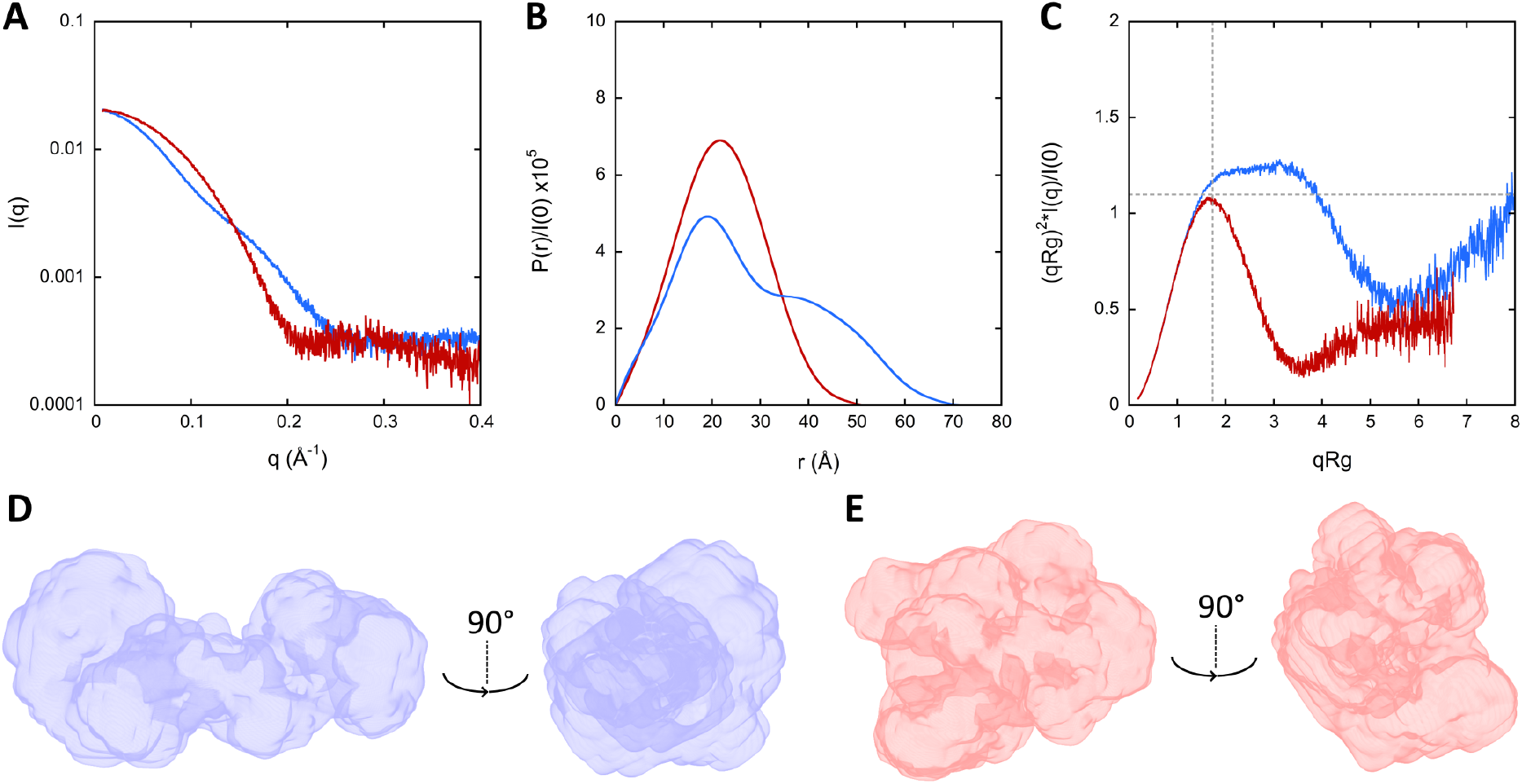
SAXS patterns and DENSS analysis of holo-CaM in the absence and in the presence of CDZ. **A**. Scattering patterns of holo-CaM (blue trace) and holo-CaM:CDZ complex (red trace). **B**. Distance distribution function P(r) of holo-CaM in the absence and in the presence of CDZ using GNOM, same color code as in panel A. **C**. Dimensionless Kratky representation of holo-CaM in the absence and in the presence of CDZ, same color code as in panel A. Dash lines crossing x-axis (√3) and y-axis (1.104) indicate the characteristic peak for globular proteins. **D. and E**. DENSS models of holo-CaM and holo-CaM:CDZ, respectively.

### 2.3. Structural analysis of holo-CaM by SAXS and X-ray crystallography

The Ensemble Optimization Methods (EOM) suite of programs^38,39^ was used to generate an ensemble of structural models of holo-CaM. Briefly, ten thousand conformations were generated by the Ranch program, starting from the structures of the two calcium-bound lobes of holo-CaM (pdb 1CLL^40^). Residues (76 – 81) in the interlobe linker region were described as dummy residues. These residues were then substituted by a full-atom description with PD2^41^ and the energy of each linker was minimized using SCWRL4^42^. The scattering pattern of each of the 10000 conformations was then computed using CRYSOL^43^. Finally, the Gajoe program was used to select ensembles of conformation, the average scattering pattern of each ensemble being fitted against the experimental data using a genetic algorithm. The best ensemble for holo-CaM (SASDNX3) displays the typical conformation of holo-CaM with two well-folded lobes connected by a highly flexible linker (Figure 2A) and shows a good fit to the experimental scattering pattern (Figure 2B) with a χ^2^ value of 1.066. The four models in this ensemble fit quite well inside the DENSS^37^ volume (Figure 2C).

**Figure 2.**
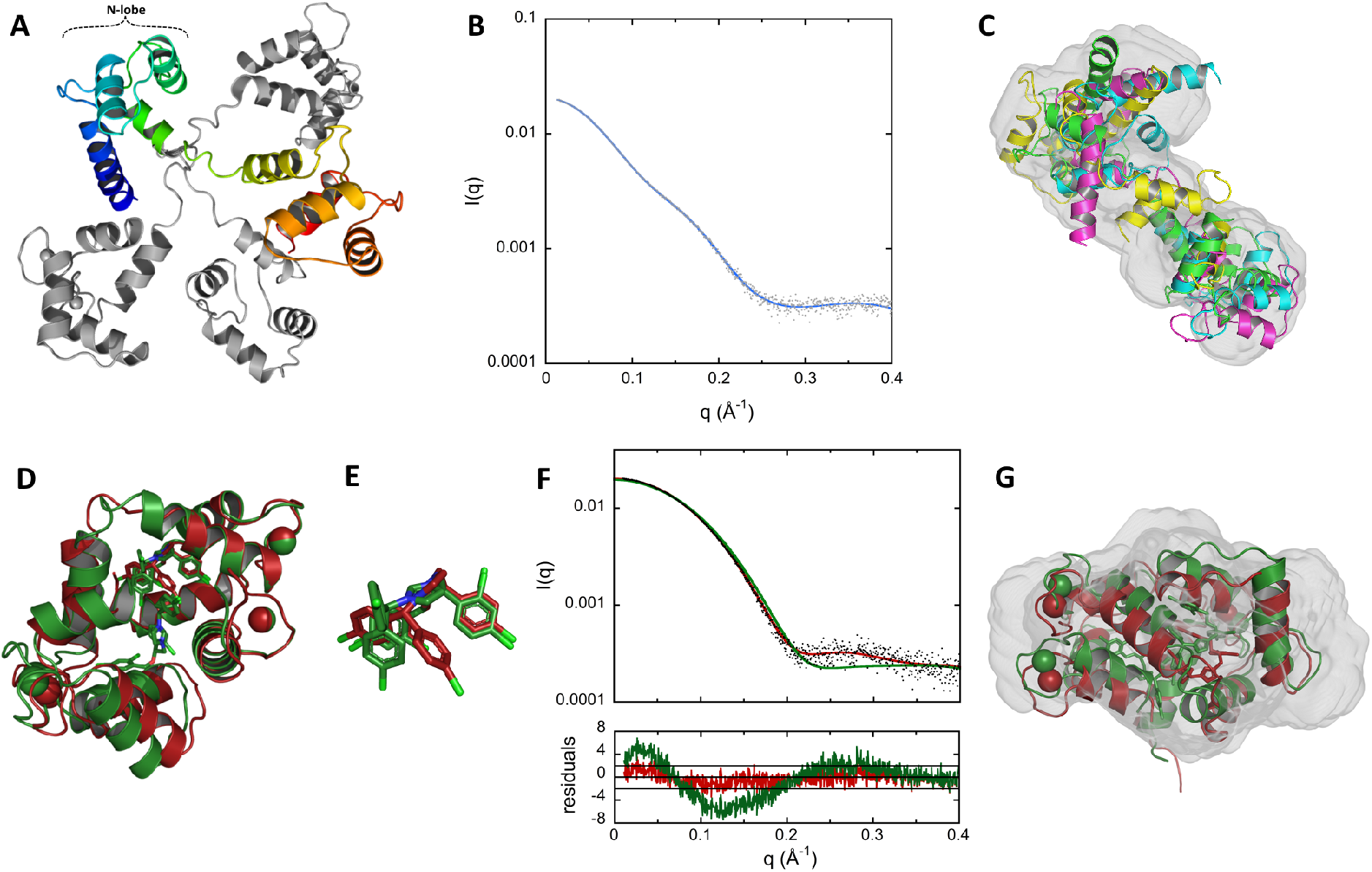
Structural models of holo-CaM using EOM and structures of holo-CaM:CDZ determined by X-ray crystallography. **A**. Final ensemble of SAXS-derived conformations of holo-CaM using EOM. The four structural models (SASDNX3) are superimposed by aligning their N-lobe alpha carbons (residues 6 to 66) using Pymol. One structural model is shown in rainbow, the three others are in grey. **B**. Comparison of experimental data (grey dots) to the calculated scattering pattern (blue curve) of the final EOM ensemble. **C**. Fitting of the four EOM structural models of holo-CaM to the SAXS-derived DENSS volume. **D**. X-ray structures of holo-CaM in complex with one (red, pdb ID: 7PSZ) and two (green, pdb ID: 7PU9) CDZ molecules. **E**. Superimposition of the CDZ molecules of both structures showing the rotation of the chlorophenyl moieties. **F**. Top: Fitting of the calculated scattering patterns of the two crystallographic structures of holo-CaM:CDZ obtained using Crysol to the experimental SAXS pattern **recorded** with 333 μM of CaM and 1050 μM of CDZ. The χ^2^ are 1.4 and 9.3 for the 1:1 (red) and 1:2 (green) holo-CaM:CDZ complexes, respectively. Bottom: Distribution of reduced residuals corresponding to the two fits **presented above**. (**G**) Fitting of the two crystallographic structures of holo-CaM:CDZ to the SAXS-derived DENSS volume (SASDNY3).

The holo-CaM:CDZ complex was investigated using X-ray crystallography. Depending on the molar ratio of holo-CaM and CDZ, we obtained two crystal forms. For the holo-CaM:CDZ complex prepared at a molar ratio of 1:2.2 (1 mM holo-CaM), the unit-cell parameters, merging statistics and systematic absences were consistent with space group C121, while for the holo-CaM:CDZ complex prepared at a molar ratio of 1:10 (1 mM holo-CaM), the parameters were coherent with space group P6_1_22. The corresponding structures pdb 7PSZ (holo-CaM:CDZ_A_ for holo-CaM with one CDZ molecule, CDZ-A) and pdb 7PU9 (holo-CaM:CDZ_BC_ for holo-CaM with two CDZ molecules, CDZ-B and CDZ-C) were solved by molecular replacement to 1.9 Å and 2.3 Å, respectively (Figure 2D and Table S2). In the complex prepared at a 1:2.2 molar ratio of holo-CaM:CDZ (pdb 7PSZ), some density was left unattributed inside holo-CaM. This density, which could be assigned to a second CDZ, may arise from partial occupation due to differences between molecules within the unit-cell or random sub-stoichiometric binding.

The two crystal structures of holo-CaM:CDZ with one and two CDZ molecules are quite similar with a RMSD of only 1.4 Å over all Cα and exhibit compact and globular structures (Figures 2D). CDZ-A and CDZ-B are bound to the same region of holo-CaM and are mostly localized in the N-lobe. The superimposition of the N-lobes of the two crystal structures shows that the conformation of CDZ-A and CDZ-B slightly differ by the rotation of two chlorophenyl groups (Figure 2E). This rotation seems required to accommodate the second CDZ (CDZ-C) inside the holo-CaM:CDZ_BC_ complex. Analysis with LigPlot+^44^ indicates that holo-CaM:CDZ interactions are mostly driven by hydrophobic effects (Figure S4), with CDZ-A and CDZ-B mainly interacting with residues from the C-terminal part of the N-lobe and the N-terminal part of the C-lobe of holo-CaM, as shown in Table S3. These residues are similar to those identified by Reid *et al*.^45^. CDZ-C mainly interacts with the N- and C-terminal extremities of CaM.

The crystal structures and the SEC-SAXS data of holo-CaM:CDZ were compared using DENSS and Crysol. Both crystal structures fit well within the envelope of the DENSS model generated with the SEC-SAXS data of the holo-CaM:CDZ complex (Figure 2G). Comparison of the theorical SAXS patterns of the two crystal structures generated by Crysol to the experimental SEC-SAXS pattern of the complex indicates that the holo-CaM:CDZ_A_ crystal structure is in excellent agreement with experimental data with a χ^2^ of 1.4, against 9.3 for the holo-CaM:CDZ_BC_ crystal structure (Figure 2F). Taken together, these analyses indicate that the holo-CaM:CDZ complex exhibits a similar conformation in solution and in the crystals and suggest that a single CDZ molecule is sufficient to induce an open-to-closed conformational change of holo-CaM. The part of CDZ_A_, which remains solvent-exposed once interacting with the N-lobe of holo-CaM, likely forces the C-lobe to collapse by a hydrophobic effect, closing the two lobes of holo-CaM on themselves, with holo-CaM wrapping around CDZ_A_.

### 2.4. HDX-MS analysis of holo-CaM upon CDZ binding

The effect of CDZ binding on the solvent accessibility of holo-CaM was also investigated by HDX-MS. The holo-CaM:CDZ complex used in HDX-MS was performed at a 1:32 molar ratio (0.63 μM CaM and 20 μM CDZ) in the presence of 2% DMSO (Table S4, Figures S5 and S6). The excess of CDZ ensures that most of holo-CaM (circa 90 %) remains complexed during labelling (*i.e*., using a K_D_ of 3 ± 2μM and a 1.2 binding stoichiometry). CDZ binding induces similar reductions in solvent accessibility within the N- and C- lobes of holo-CaM (Figures 3A, 3B and 3C), and has no effect on regions covering residues 2-9 (N-ter), 73-84 (interlobe linker region) and 114-124 (helix α6). The effects of CDZ are best illustrated and visualized in Figure 3D. This figure was generated by plotting on the CaM:CDZ_A_ crystal structure (pdb 7PSZ, Figure 2D) the average “Differences in Uptake differences” calculated between CDZ-bound and free holo-CaM for all labelling time points (Figure S6). The main reductions in solvent accessibility are located in region 21-45 (calcium binding loop 1, α2) of the N-lobe, and in regions 84-90 (α4b), 104-113 (α5) and 143-149 (α7) of the C-lobe. The regions with high reductions in solvent accessibility identified by HDX-MS compare well with the residues interacting with CDZ-A and CDZ-B in the crystal structures, as highlighted with Ligplot+ (Table S3). The residues interacting with CDZ-C in the crystal structure experience only a weak reduction in solvent accessibility, suggesting that CDZ-C is essentially absent in holo-CaM:CDZ complexes at the concentrations used for HDX-MS experiments. Finally, a weak reduction in solvent accessibility is measured for the interlobe linker region. The HDX-MS results appear therefore consistent with the crystal structure of the holo-CaM:CDZ_A_ complex and confirm that CDZ binding to the N-lobe dramatically stabilizes both lobes.

**Figure 3.**
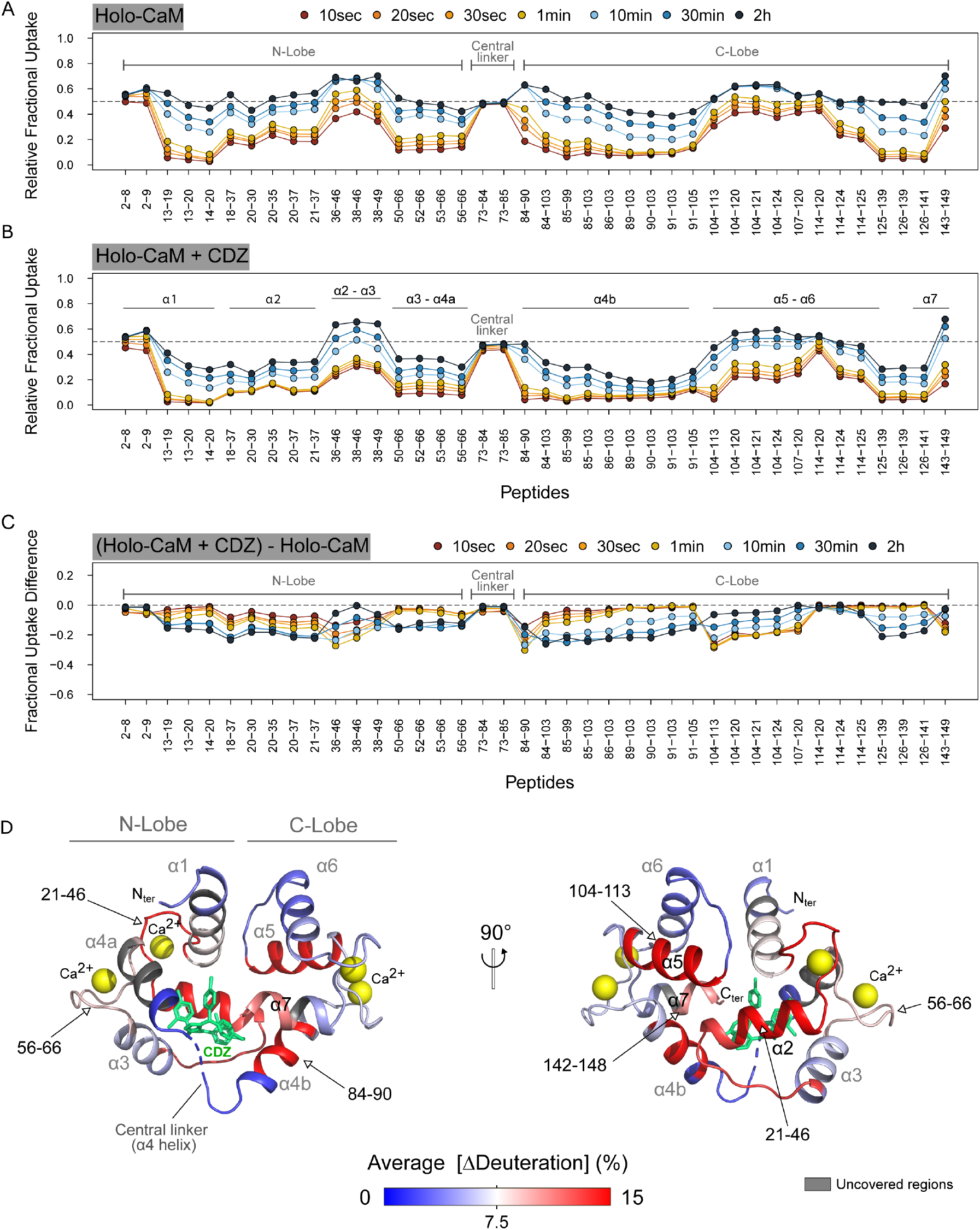
Effects of CDZ binding on the deuterium uptake profile of holo-CaM. **A. and B**. Relative fractional uptake plots of holo-CaM measured in the presence and in the absence of 20 μM CDZ. Each dot corresponds to the average uptake value measured in three independent replicates. **C**. The effects of CDZ binding on holo-CaM are visualized on the Fractional Uptake Difference plot. Negative values indicate a reduction in solvent accessibility induced by CDZ binding. **D**. Cartoon representation of holo-CaM showing the average differences in “Fractional Uptake Differences” between the CDZ-bound and free holo-CaM states. The Fractional Uptake Differences ([ΔDeuteration] in %) measured between the CDZ-bound and free states were extracted for each peptide at each labelling time point, averaged, and plotted on the crystal structure of CaM:CDZ_A_ (pdb 7PSZ). CDZ-A is colored in green. The average ΔDeuteration values [Average (ΔDeuteration)] are colored from blue (no variation) to red (major reductions in uptake).

The effects of CDZ binding on the exchange behavior of holo-CaM were finally compared to those induced by the binding of CaM-binding peptides: two peptides, the H-helix and P454 derived from the adenylate cyclase toxin CyaA from *Bordetella pertussis*^46–49^ and MLCK peptides.^10,49,35^ The interlobe linker region (peptide 73-84) remains accessible to the solvent in the absence and in the presence of ligands (Figure S7). Interestingly, CDZ, the MLCK and the H-helix peptides induce similar differences in HDX uptake of holo-CaM (albeit with distinct amplitudes but the global patterns remain very similar). These results indicate that CDZ mimics the effects of biological ligands on holo-CaM. Taken together, HDX-MS data shows that CDZ binding to holo-CaM dramatically reduces the solvent accessibility of both lobes without affecting the interlobe linker region.

### 2.5. Effect of CDZ binding on holo-CaM hydrodynamic parameters monitored by NMR

We evaluated the influence of CDZ binding to holo-CaM on the tumbling correlation time (τ_c_) at 37 °C with increasing equivalents of CDZ (Table S5). The observed increase in τ_c_ indicates that while the N- and C-lobes of holo-CaM tumble independently of each other (τ_c_ = 4.5 ± 0.2 ns), the two lobes tumble together in a more compact holo-CaM:CDZ 1:1 complex (5.2 ± 0.3 ns). The τ_c_ value of CDZ-bound holo-CaM agrees well with the value calculated for a globular, compact protein of molecular mass and partial specific volume corresponding to holo-CaM (5.2 ns), further confirming the SAXS and X-ray analyses.

### 2.6. Effects of CDZ on holo-CaM NMR spectra and chemical shifts

We recorded ^1^H-^15^N correlation spectra of holo-CaM in the absence and in the presence of 0.5 and 1 equivalents of CDZ (Figure 4A). CDZ binding dramatically perturbs most holo-CaM signals, indicating that CDZ affects not only the signals of residues at the binding interface but also exhibits long-range effects on the conformations and/or internal motions throughout most of the protein. The spectrum of holo-CaM with a half-equivalent of CDZ (cyan, Figure4A, right panels) corresponds to the sum of the spectra of the free and the 1:1 molar ratio of holo-CaM:CDZ complex. Thus, (i) the free and bound holo-CaM conformations are in slow exchange on the chemical shift time scale as expected for a high affinity interaction and (ii), at sub-stoichiometric concentrations, one CDZ molecule binds to one holo-CaM. Binding of CDZ also affects the relative intensities of the ^1^H-^15^N signals and hence the internal motions of holo-CaM (see 130I signal in Figure 4A).

**Figure 4.**
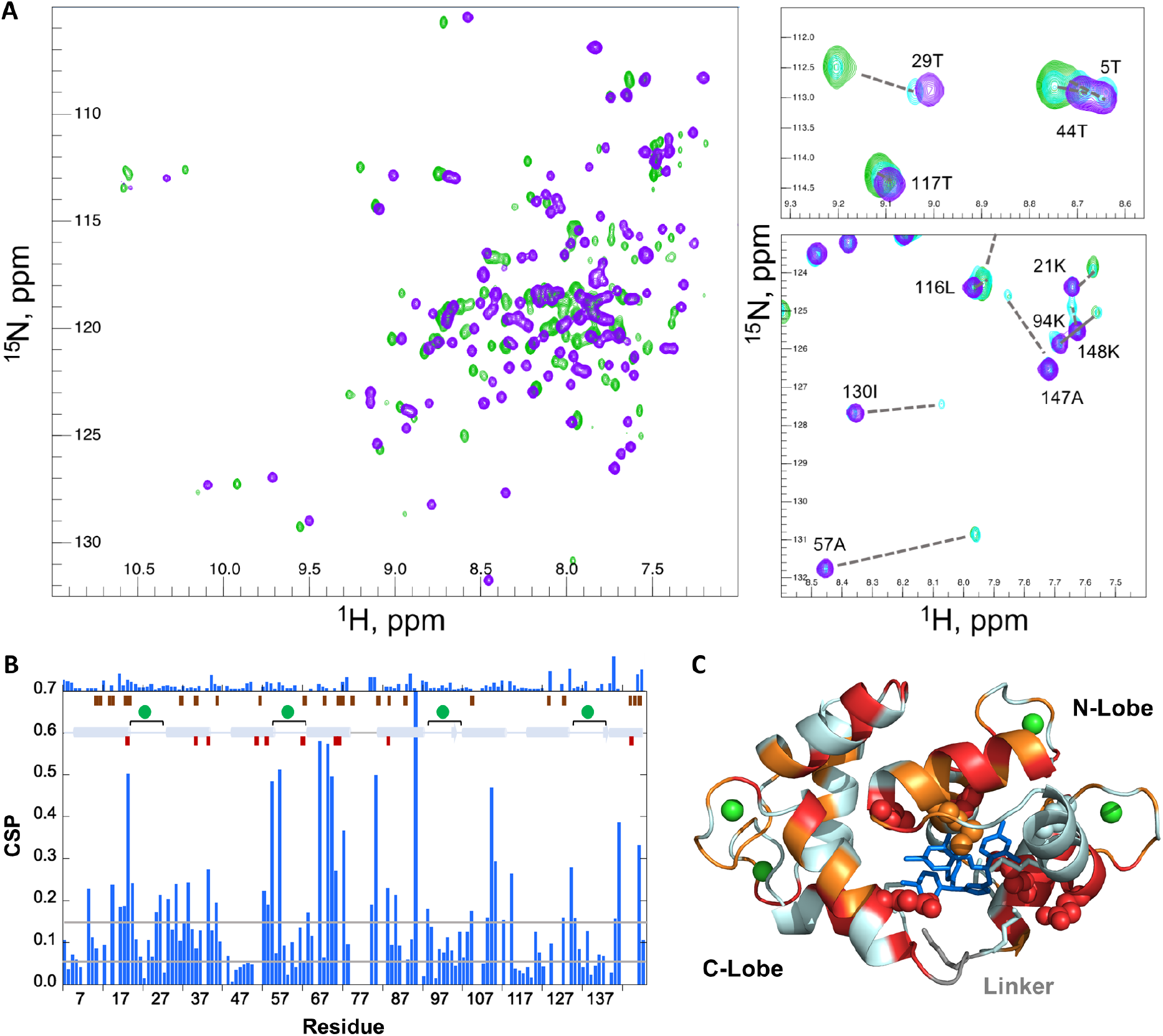
CDZ binding monitored by CaM ^1^H-^15^N chemical shift perturbation (CSP). **A**. (left) ^1^H-^15^N SOFAST full fingerprint spectra (recorded at 37 °C) of holo-CaM alone (mauve) and in the presence of 1.0 equivalent of CDZ (green). (right) Zoom on two selected regions of the fingerprint spectra. The spectrum of holo-CaM in the presence of 0.5 equivalents (cyan) is also displayed. The assignments of free holo-CaM are shown and the dotted lines indicate the corresponding signal in the 1:1 holo-CaM:CDZ sample. **B**. CSP values of 1 CDZ equivalent added to holo-CaM as a function of the residue number. The secondary structure (helix=cylinder, strand=arrow), calcium binding loops (spheres and square brackets) and linker region (grey line) are schematized. The CSP values between the 1:1 and 1:2 holo-CaM:CDZ complexes are displayed on the top of the panel, respecting the same scale. The positions of contacting residues in the X-ray 1:1 and 1:2 holo-CaM:CDZ complex structures are represented by wine and maroon rectangles, respectively. Grey lines represent the CSP values chosen as thresholds for very strong and strong CSPs. **C**. Residues with very strongly perturbed (red, CSP ≥ 0.14) and strongly perturbed (orange, 0.07 ≤ CSP < 0.14) amide resonances are highlighted on the cartoon representation of the 1:1 holo-CaM:CDZ x-ray complex structure. CDZ is shown as blue sticks and Ca^2+^ ions as green spheres. The side chains of the CaM residues in close contact with CDZ as defined by Ligplot+ are highlighted as spheres if assigned (19, 36, 39, 54, 63, 71, 84) or as sticks (51, 72, 76, 77 and 145) if not observed by NMR. Amide resonances of the following residues were not assigned in the holo-CaM:CDZ complexes: 8, 12, 14, 16, 38, 51-52, 72, 75-79 ,82-83 ,88, 92, 106-107, 112, 114, 124, 126-127, 129-130, 139, 143-146.

As evaluated from the backbone and CB resonances of the free and CDZ-bound holo-CaM using Talos-N, CDZ binding did not alter the secondary structure of holo-CaM, which was consistent with the secondary structure of the crystal structures determined herein and of holo-CaM in solution^50^ (Table S6). Hence, the variations of chemical shifts in ^1^H-^15^N correlation spectra were due to residues in contact with CDZ, and/or to a modification of the tertiary structure and/or the internal dynamics of the protein.

We then performed a quantitative analysis of the chemical shift perturbations (CSPs) of holo-CaM upon binding to CDZ (Figures 4B and 4C). Most of the highly affected residues were localized in regions contacting CDZ. Some strong perturbations were also located far away from the binding interface, for instance in calcium binding loops. Overall, the CSPs are in agreement with the binding interface observed in the holo-CaM:CDZ_A_ complex (Figure 4C) and further indicate that CDZ binding affects distal regions in holo-CaM. Importantly, of the 31 unassigned amide resonances (mainly because of exchange broadening due to conformational exchange on the μs-ms time range), 18 were localized in the C-lobe, 5 on the linker region and neighboring residues and only 8 in the N-lobe, suggesting that exchanges between different conformations in the complex take place on the μs-ms time range, particularly affecting the C-lobe, which establishes less contacts with CDZ, as well as the linker region. In holo-CaM, the backbone and CB chemical shifts of residues 77-81 of the interlobe linker are consistent with a highly dynamic segment, as indicated from chemical shift-derived S^2^ order parameters determined using RCI (Figure S8). However, in the presence of CDZ, we observed a shift and exchange broadening of the corresponding amide signals (as well as those of neighboring residue signals 75-79 and 82-83), with residues 80D, 81S, 84E and 74R displaying very strong CSPs (Figure 4B).

Addition of an excess of CDZ to holo-CaM brings only minor modifications to the spectrum of holo-CaM bound to one CDZ, as evidenced by the very low CSP values observed between the holo-CaM:CDZ complexes at a molar ratio of 1:1 and 1:2.9 (Figure 4B). This relatively small effect, that might reflect binding of a second CDZ molecule, can be rationalized considering that (i) the binding sites and residues in contact with CDZ are similar in both complexes for CDZ_A_ and CDZ_B_, (ii) several residue signals close to CDZ_C_ were exchange broadened and could not be assigned (see Figure 4 legend for details) and (iii) the structures of one- and two-CDZ loaded holo-CaM are similar.

Whereas binding of CDZ to holo-CaM is a slow process in the chemical shift time scale, 1H-^15^N spectra indicate that some residues undergo exchange between different conformations on a faster time scale within the complex. This phenomenon, which gives rise to small chemical shift variations that depend on CDZ concentration, implicates residues in the 2^nd^, 3^rd^ and 4^th^ calcium binding loops, with residues such as 57A (2^nd^ loop) and 137N (4^th^ loop) showing the highest effect.

### 2.7. Influence of CDZ on the internal dynamics of holo-CaM

The internal dynamics of free and CDZ-bound holo-CaM were compared using the ^15^N transverse (T_2_), longitudinal (T_1_) and heteronuclear ^1^H-^15^N nOe relaxation parameters (Figures S9 and S10). In holo-CaM, the relaxation parameters (high T_2_, low T_1_, low T_1_/T_2_ and low ^1^H-^15^N nOes) of residues in the linker region (77-81) and flanking residues in α-helices H4 and H6 are characteristic of high amplitude motions on the ns-ps time scale, with a higher flexibility around its mid-point (T79-D80). In contrast, the relaxation parameters of the α-helical elements of the N- and C-lobes are indicative of ordered regions in a globular protein. Binding of CDZ drastically changes the internal dynamics of holo-CaM.

A simple picture of the modification of the amplitude of the fast motions caused by CDZ binding can be obtained from the difference of the nOe values between the bound and free conformations (Figure 5). Most residues with available data displayed higher nOes within the complex, with 44 residues (highlighted in red and orange in Figure 5) showing a significant CDZ-induced effect. These data suggest a global reduction of internal fast motions of holo-CaM within the complex. Albeit residues in all the secondary structure elements and calcium binding loops showed this increase in nOe, the latter was not evenly distributed throughout the structure. Among calcium binding loops, loop 3 had the higher increase in nOe. This stabilizing effect was less pronounced for α-helix H1, which is far away from the CDZ binding site. Importantly, the increase of nOe for residues 80D and 81S in the linker was of 0.4, which indicates that CDZ binding restricts the fast motions in the linker. The fact that residues 77-79 in the linker and flanking residues 75-76 (unassigned) were highly impacted by CDZ binding and exchange-broadened, further shows that the linker experiences conformational modifications and motion restrictions also on a slower time scale (μs-ms). This motion restriction is however modest and the linker region remains dynamic in the complex as evidenced by HDX-MS that probes dynamics on a longer time scale (< minutes). As observed with CSPs, binding of a second CDZ molecule (sample with a three-fold excess of CDZ) resulted in modest modifications in the ^15^N T_1_, T_2_ and ^1^H-^15^N nOe profiles of holo-CaM relative to the 1:1 complex (Figure S10). No clear evidence of different rigidity in the 1:1 or 1:2 complexes was detected. Taken together, the sub-nanosecond internal dynamics of holo-CaM:CDZ 1:1 and 1:2 complexes are similar (and reduced relative to the free protein), and holo-CaM experiences slow μs-ms conformational changes within the complex.

**Figure 5.**
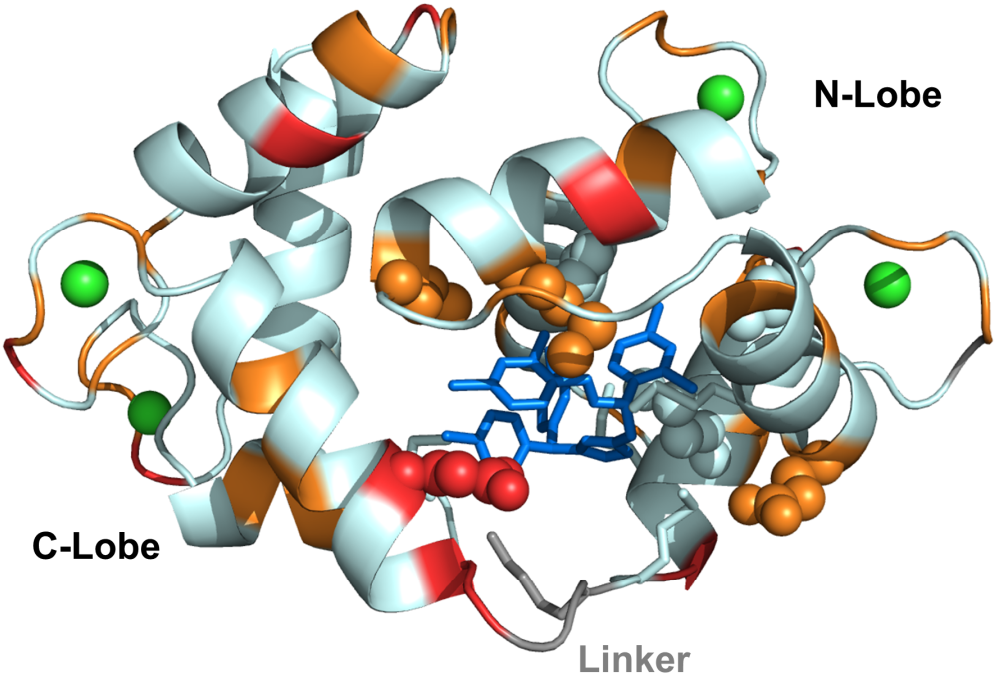
Decrease of backbone amide internal motion amplitudes in the nanosecond-picosecond time scale of holo-CaM upon binding to CDZ. The difference of the heteronuclear ^1^H-^15^N nOe of CDZ bound and free holo-CaM is color-coded on the cartoon representation of the holo-CaM:CDZ 1:1 X-ray structure. Red indicates very high positive nOe differences (≥ 0.18) and orange high nOe differences (between 0.10 and 0.16), denoting a decrease in the amplitude of fast internal motions (ns-ps) in the complex. CDZ is shown as blue sticks, Ca^2+^ ions are displayed as green spheres and the linker residues are shown in grey. The side chains of the CaM residues in close contact with CDZ as defined by Ligplot+ are highlighted as spheres if assigned (19, 36, 39, 54, 63, 71, 84) or as sticks (51, 72, 76, 77 and 145) if not observed by NMR.

## 3. Conclusion

In the present work we have explored the structural basis of holo-CaM interaction with CDZ, one of the most potent CaM antagonists known to date. CDZ has been widely used to probe the role of holo-CaM in various biological processes.^52,33,22,34^ How this molecule interacts with holo-CaM and inhibits its regulatory activity had not been elucidated at the molecular level. Although numerous studies have been carried out to define the biochemical and biophysical parameters of holo-CaM:CDZ interaction, major uncertainties remained regarding the structure, dynamics, affinity, and stoichiometry of the complex.

Our present results reveal a main CDZ binding site involving primarily the N-lobe of CaM, although a second CDZ could be detected in crystals grown at high CDZ:holo-CaM ratio and high concentration (10 mM CDZ). Solution structure analysis indicated only a very low proportion of holo-CaM complexed with two CDZ (holo-CaM:CDZ_BC_), which is unlikely to be of any relevance in physiological conditions, *i.e*., when CDZ is used to probe the potential implication of CaM in signaling pathways and/or physiological processes (usually at concentrations below 50 μM). Earlier work, mainly based on indirect measurements using fluorescent probes, had speculated on multiple CDZ binding sites - up to six molecules per CaM^33^, but our data clearly rule out such models.

Structures of other CaM:inhibitor complexes previously revealed that CaM may accommodate a variable number of antagonists in distinct calcium-loaded conformations, (*e.g*. 2 or 4 TFP per CaM or 2 arylalkylamine antagonists, etc.), and even two distinct drugs simultaneously for example, trifluoperazine and KAR-2 or vinblastine and KAR-2, but not trifluoperazine and vinblastine^26^. This diversity may reflect the somehow artificial conditions of crystallization that use high concentrations of compounds (and CaM) and are likely favoring hydrophobic forces between drugs and the hydrophobic cavities of CaM, a consideration that should be kept in mind when interpreting the corresponding crystal structures.

Our integrative structural biology approach indicates that CDZ binding dramatically affects the conformational dynamics of CaM that collapses from a dumbbell-shaped conformation into a compact globular structure. Despite the fact that CDZ mainly contacts the N-lobe, it triggers a complete closing of the two lobes of holo-CaM into a structure that appears very similar to that of CaM bound to classical CBS peptides (*e.g*. MLCK^53,51,49^). The CDZ-induced compaction was clearly established both in crystal and in solution. The CDZ-induced stabilization of this closed conformation thus locks CaM into an inactive form unable to associate with most CBS of target proteins. Even a target enzyme, such as *B. pertussis* CyaA that is activated by interacting with the CaM C-terminal lobe only,^54,55,49^ can be efficiently inhibited by CDZ.^35^ This suggests that CDZ, by stabilizing the closed CaM globular conformation, should be capable of blunting all major CaM associations with most enzyme targets. Interestingly, this mode of binding is similar to that of TFP except that this drug, in contrast to CDZ, mainly contacts residues in the C-lobe of CaM.^25^ Hence, in both cases, drug binding, primarily to one of the two lobes of CaM, is sufficient to trigger drastic open-to-closed conformational changes of the protein.

The two crystal structures of holo-CaM:CDZ with one and two CDZ molecules revealed that the key CDZ binding residues, mainly localized within the N-lobe of holo-CaM, are mostly hydrophobic and include several methionines. Interestingly, in the 1:2 complex, one of the two CDZ molecules (CDZ-B) adopts essentially the same binding mode as the single CDZ (CDZ-A) in the 1:1 complex, although with a slightly different arrangement that is likely required for accommodating the second CDZ. NMR studies clearly indicated that, within the 1:1 holo-CaM-CDZ complex, some residues undergo exchange between different conformations on a fast (μs-ms) time scale. It is thus possible that in solution, CDZ is continuously sampling different configurations within the closed holo-CaM-CDZ globular complex, a feature that was also recently observed for another antagonist idoxifene.^30^ Binding of a second CDZ molecule mainly onto the CaM C-lobe, actually triggers very little changes on the overall conformation of the complex and its internal dynamics. Structural analysis of holo-CaM-CDZ in solution is clearly in favor of a 1:1 stoichiometric complex. Overall, binding of a second CDZ is unlikely to occur in solution in standard physiological assays *in vitro* or in cells (with CDZ below 20 μM) and it is therefore reasonable to assume that in these tests, binding of a single CDZ can fully block CaM activity. This is important to consider when experimentally probing CaM signaling function with CDZ at a concentration able to match the total CaM concentration in cell (estimated to range from 3 to 10 μM, depending on cell types^35^).

Collectively, these data highlight the remarkable plasticity of CaM that can adopt highly diverse conformations to precisely fit to widely different antagonist scaffolds as well as to peptidic-binding motifs on target protein effectors. Our present results also open up new opportunities to design CDZ derivatives with substantially higher CaM-affinity, and consequently, better selectivity. Indeed, CDZ is known to affect several cellular targets, including L-type Ca^2+^, K^+^, Na^+^ channels, and sarcoplasmic reticulum (SR) calcium-release channels.^52^ At high concentrations, it may exhibit other pharmacologic effects as well. CDZ is cytotoxic at high concentrations and has been shown to induce apoptosis in breast cancer and hepatoma cells,^56,2^ to inhibit growth of murine embryonal carcinoma cells^57^ and to enhance differentiation of colon cancer cells as well.^58,2^ These adverse effects somehow limit the application of this molecule (as well as that of all other CaM antagonists thus far) in dissecting calcium/CaM signaling. A more selective targeting of CDZ analogs onto holo-CaM could restrict the off-target effects and improve its biological pertinence in fundamental research but also potentially in therapeutic applications.^59–61^ Renewed efforts are currently applied to develop improved CaM-antagonists as illustrated by Okutachi *et al*. who recently described the development of a new covalent CaM inhibitor, called Calmirasone1, to explore the cancer cell biology of K-Ras and CaM associated stemness activities.^62^ Such CaM inhibitors might find straightforward medical applications, as illustrated by Taylor and colleagues,^63^ who provided evidence that CaM inhibitors could be efficacious in rescuing a genetic disease (Diamond-Blackfan anemia, DBA) that results from increased expression of tumor suppressor. This study suggests that CaM inhibition may offer a potential therapeutic path for treatment of DBA and other diseases characterized by aberrant p53 activity.^63,64^

Taken together, the dynamics and structural data presented herein will be instrumental to design the next generation of CDZ analogs with improved selectivity for CaM and restricted off-target effects.

## 4. Experimental Section/Methods

Material and methods are described in the supplementary information file.

## Supporting Information

Supporting Information is available from the Wiley Online Library. The file contains the materials and methods section, Figures S1 to S11, Tables S1 to S6, and the supplementary references.

## Acknowledgments

C.L. was supported by Institut Pasteur (PTR 166-19) and ANR (ANR 21-CE11-0014-01-TransCyaA). I.P. was supported by the ANR grant ANR-16-CE110020-01. M.S. was supported by the Pasteur - Paris University (PPU) International PhD Program. N.C. was supported by Institut Pasteur (DARRI-Emergence S-PI15006-12B). We thank the staff of the Crystallography core facility at the Institut Pasteur for carrying out robot-driven crystallization screenings. We thank SOLEIL and ESRF for provision of synchrotron radiation facilities. We thank the staffs of the DISCO, PROXIMA-1, PROXIMA-2 and SWING beamlines for constant support and help during data collection at Synchrotron SOLEIL (St Aubin, France) and MASSIF at Synchrotron ESRF (Grenoble, France) beamlines for assistance during the X-ray diffraction data collection.

## Data Availability

All relevant HDX-MS, X-ray and SAXS data are available in supporting information. The crystal structures have been deposited on the PDB with the access codes 6YNU and 6YNS. The structural models and experimental SAXS data have been deposited on SASBDB (Small Angle Scattering Biological Data Bank, http://www.sasbdb.org/aboutSASBDB/) under the SAS codes SASDNX3 (Calcium-bound Calmodulin, including structural models) and SASDNY3 (Calcium-bound Calmodulin complexed with Calmidazolium).

## Abbreviations

CaM: calmodulin
C-lobe: C-terminal domain of CaM
CDZ: calmidazolium
HDX-MS: hydrogen/deuterium exchange mass spectrometry
holo-CaM: calcium-loaded calmodulin
MEMHDX: Mixed-Effects Model for HDX experiments
MLCK: myosin light chain kinase
MS: mass spectrometry
N-lobe: N-terminal domain of CaM
NMR: nuclear magnetic resonance
pdb: Protein Data Bank
SASBDB: Small Angle Scattering Biological Data Bank
SAXS: small-angle X-ray scattering
SEC: size exclusion chromatography.

