## supporting information for "Dynamics and structural changes of calmodulin upon interaction with its potent antagonist calmidazolium"

**Materiel and methods**

**Chemicals**

Calmidazolium Chloride (CDZ, CAS 57265-65-3, compound R_24571, M = 687.70 g/mol) was purchased from Merck (product reference 208665). CDZ was resuspended in 100% DMSO at a final concentration of 14.5 mM. CDZ was resuspended at 145 mM in 100% DMSO for the crystallography experiments of the Holo-CaM:CDZ complex prepared at a molar ratio of 1:2. The MLCK, H-helix and P454 peptides^1–3^ were synthesized by Genosphere (Paris) and prepared as previously described^1^.

**Buffers**

Experiments were performed using the following buffers:

Buffer A: 20 mM HEPES, 150 mM NaCl pH 7.4

Buffer B: 20 mM HEPES, 150 mM NaCl, 2 mM CaCl_2_, pH 7.4

Buffer C: 20 mM HEPES, 150 mM NaCl, 2 mM EDTA, pH 7.4

Buffer D: 20 mM HEPES, 150 mM NaCl, 2 mM CaCl_2_, 2% DMSO, pD 7.4 prepared in 99.98 atom % D deuterium oxide

Buffer E: 20 mM HEPES, 150 mM NaCl, 2 mM CaCl_2_, 2% DMSO and 20 μM CDZ, pD 7.4 prepared in 99.98 atom % D deuterium oxide

Buffer F: 20 mM HEPES, 150 mM NaCl, 2 mM CaCl_2_, 5% DMSO, pH 7.4

**Calmodulin purification**

Human Calmodulin (CaM) was produced in *E. coli* and purified as previously described^2,4,5^. Briefly, CaM was precipitated with ammonium sulfate followed by a glacial acetic acid precipitation. Then, CaM was purified as follows: a first HIC on Phenyl Sepharose (Apo-CaM) in the presence of EDTA, an IEC on Q-Sepharose fast flow, a second HIC on Phenyl Sepharose (Holo-CaM) in the presence of calcium (and eluted in EDTA) and a SEC on Sephacryl S100 equilibrated in buffer A. Calmodulin is stocked in buffer A. Protein concentration was determined by spectrophotometry (ε_280nm_ = 2,980 M^-1^ cm^-1^) and protein integrity was checked by SDS-PAGE and intact mass spectrometry analysis (M = 16706.03 g.mol^-1^) by ESI-Q-ToF. The MS analysis indicates that the N-terminal methionine residue is post-translationally cleaved in *E. coli*.

**Isothermal titration calorimetry**

ITC experiments were performed using a VP-ITC calorimeter (Malvern Panalytical, Orsay, France). The ITC experiments were performed in buffer F at 25°C (unless otherwise stated). For a typical titration, the solution of analyte (8 µM of Holo-CaM) was loaded in the reaction cell. The titrant (200 µM of CDZ) was loaded into the syringe before injection of 5 to 15 μl of the titrant into the reaction cell at intervals of 600 seconds. Heats of dilutions were measured by injecting the titrant into the buffer and were subtracted from the heat of reaction. The titration profiles were analyzed using the Origin7 software (OriginLab, Northampton, MA, USA) to determine the thermodynamic parameters, as previously described ^3^.

**Synchrotron Radiation Circular Dichroism**

Synchrotron radiation circular dichroism (SRCD) was performed on the DISCO beamline of the synchrotron SOLEIL (Saint-Aubin, France). Spectra were recorded at 25°C with an integration time of 1.2 s and a bandwidth of 1 nm with a resolution of 1 nm. A far-UV spectrum represents the average of four individual scans. QS cells (Hellma, France) with a pathlength of 20, 50, 100 or 200 μm (depending on final protein concentration) were used to record spectra in the far-UV range (from 190 to 250 nm). CD spectra were measured in buffers A and B. The addition of DMSO, due to the use of CDZ resuspended in 100% DMSO, did not exceed 0.9% of DMSO in samples for SRCD. The CD unit is the mean residue ellipticity, MRE, expressed in (kilodegrees*cm^2^) / (dmol*residues), and calculated as previously described ^5,6^.

**Small Angle X-Ray Scattering**

X-ray scattering data were collected at the SWING beamline^7^ of the SOLEIL Synchrotron (Saint-Aubin, France) (Table S1 gives all experimental details in accordance with BioSAXS publication guidelines^8^). Measurements were performed using a size-exclusion HPLC column (TSKgel G3000SW) online with the SAXS measuring cell, a 1.5 mm diameter quartz capillary contained in an evacuated vessel ^9^. All sample were prepared in buffer B with 5-7% final DMSO (depending on CDZ concentration). Briefly, 50 μL of sample solution were loaded onto the equilibrated column. Scattering of the elution buffer before void volume was recorded and used as buffer scattering for subtraction from all protein patterns. Successive frames of 1 s were recorded. The elution flow of 0.5 mL/min ensured that no protein was irradiated for more than 0.4 s. Primary data reduction was performed using Foxtrot, the SWING in-house software. This yielded azimuthally averaged scattering intensities I(q) put on absolute scale (cm^-1^ units) using water scattering, where q is the momentum transfer (q = 4π sinθ/λ, where 2θ is the scattering angle and λ the wavelength of the X-rays). Data was subsequently processed using the program package PRIMUS ^10^. The forward scattering I(0) and the radius of gyration (Rg) were evaluated using the Guinier approximation ^11^. Frames over the elution peak were analyzed individually before averaging the appropriate subset that yielded identical I(q)/c profiles with US-SOMO^12^. The distance distribution function P(r) was determined using the indirect Fourier transform method as implemented in the program GNOM ^13^. The molecular mass was estimated from the value of the forward scattering. The program DENSS ^14^ was used to calculate electron density maps directly from scattering curves.

**Modelling Holo-CaM using EOM**

We undertook Holo-Cam modeling in terms of ensembles of conformations using the package EOM (Ensemble Optimization Method) ^15,16^. EOM is a well-suited program suite to describe a flexible protein such as Holo-CaM. The program Ranch within EOM creates a large (10,000) pool of random conformations using a chain of dummy residues describing the protein. The program offers a choice between three chain types to be generated corresponding to the use of three Cα angle distributions: random coil, native and compact. The native option was used. On average, random models will be more extended than native-like, while the “compact” option will force the reconstructed linkers to be rather compact. Dummy residues are subsequently substituted by complete residues using the programs PD2 ^17^ and SCWRL4 ^18^ before calculating all scattering patterns using Crysol ^19^. The routine Gajoe within EOM tries to fit the experimental scattering curve by the average of the calculated scattering patterns of an ensemble of conformations using a genetic algorithm protocol. The program is run many times and yields equally good fits with different ensembles. The resulting conformations in any ensemble should therefore be considered as illustrations of the polypeptide chain main features, rather than actual conformations adopted by the protein.

**Crystallization of Holo-CaM:CDZ complexes**

The crystallization experiments were performed at 18°C by the sitting drop vapor diffusion technique in 96-well plates, according to established protocols at the Crystallography Core Facility of the Institut Pasteur ^20^. For the initial screenings, sitting drops of 400 nL (1:1 protein to precipitant ratio) were set up in 96-well Greiner plates with a Mosquito automated nanoliter dispensing system (TTP Labtech, Melbourn, UK). The plates were then stored in a RockImager (Formulatrix, Bedford, USA) automated imaging system to monitor crystal growth. Initial crystallization hits were manually optimized by the hanging drop vapor diffusion technique in 24-well plates at 18 °C. The best crystals of the Holo-CaM:CDZ complex were obtained by co-crystallization in hanging drops containing 1 mM of Holo-CaM and 2.2 mM of CDZ in buffer B mixed with 2.8% DMSO, 30 %w/v PEG 8K and 0.2 M (NH_4_)_2_SO_4_ in the reservoir for the “1:1” complex and 1 mM of Holo-CaM and 10.7 mM of CDZ in buffer B mixed with 7.4% DMSO, 0.2 M CaCl_2_, 0.1 M Tris pH 8.5 and 25 %w/v PEG 4K in the reservoir for the “1:2” complex.

**Diffraction data collection and structure determination**

Diffraction data were collected at beamlines PROXIMA 1 and PROXIMA 2A (synchrotron SOLEIL, St. Aubin, France) and processed with autoPROC ^21^. The crystal structures of the complexes were solved by the molecular replacement method with Phaser ^22^, using separately the N-ter and C-ter lobes of Holo-CaM as search models (both from the PDB entry 1CTR). Restrains for CDZ were generated using the Grade web server (<http://grade.globalphasing.org/>). Final models of the Holo-CaM:CDZ complexes were obtained through interactive cycles of manual model building with Coot ^23^ and reciprocal space refinement with Buster ^24^. Polder maps ^25^ were calculated using Phenix ^26^. X-ray diffraction data collection and models refinement statistics are summarized in Table S2.

**Accession codes**

Atomic coordinates and structure factors for the Holo-CaM:CDZ complexes have been deposited in the RCSB Protein Data Bank under the accession code 7PSZ and 7PU9.

The molecular model and experimental SAXS data have been deposited on SASBDB (Small Angle Scattering Biological Data Bank, http://www.sasbdb.org/aboutSASBDB/) under the SAS codes SASDNX3 (Calcium-bound Calmodulin, including structural models) and SASDNY3 (Calcium-bound Calmodulin complexed with Calmidazolium).

**HDX-MS experiments**

**Biological sample and chemical for HDX-MS**

The initial CaM stock solution was at 276 μM in buffer A. The Holo-CaM working solution was prepared at 10 μM in buffer B and incubated for 30 min at room temperature before use. Initial stock solution of Calmidazolium (CDZ) at 14.5 mM in 100% DMSO. Working solution prepared at 1 mM in 100% DMSO and stored at -20°C.

**Sample preparation for HDX-MS**

A summary of the HDX data is provided in supplementary Table S4, following recommendations for HDX-MS reporting ^27^. The labelling was performed at room temperature using two distinct deuterated buffers (D and E) prepared in 99.98% deuterium oxide.

The Holo-CaM:CDZ complex was formed by mixing 9.4 μL of Holo-CaM (10 μM) with 0.6 μL of CDZ at 1 mM in a final volume of 30 μL buffer B. A CDZ-free Holo-CaM control was prepared in parallel by replacing CDZ by DMSO. After 1 h incubation at room temperature, continuous labelling was initiated by adding 120 µL of deuterated buffer to 30 µL of each equilibrated protein solution (i.e., CDZ-free Holo-CaM with Buffer D, CDZ-bound Holo-CaM with Buffer E). The final deuterium excess was ~80% to favor unidirectional exchange. The DMSO concentration was maintained at 2% during labelling. The exchange reaction was quenched after 10 sec, 20 sec, 30 sec, 1 min, 10 min, 30 min and 120 min by mixing 20 µL of the labelling reaction (i.e., 12.54 pmol of Holo-CaM) with 40 µL of quench buffer (2.5% formic acid, 4M urea) maintained at 4°C to achieve a final pH of 2.5 (final D_2_0/H_2_0 ratio: 0.27/0.73). Quenched samples were immediately snap-frozen in liquid nitrogen and stored at -80 °C. Undeuterated controls were treated using an identical procedure. Triplicate labelling experiments were performed for each time point and condition for all HDX-MS analyses (independent technical replicates).

**HDX-MS data acquisition**

HDX-MS analyses were performed with the aid of an HDX manager (Waters Corporation, Miliford, MA) maintained at 0 °C. Prior to mass analysis, samples were rapidly thawed and 10.45 pmol (i.e., 50 μL) of labelled Holo- or Apo- CaM (with or without CDZ) were digested for 2 min at 20 °C using an in-house packed immobilized pepsin column (2.0 x 20 mm, 66 µL bead volume; immobilized pepsin from Thermo Scientific). Peptides were trapped, concentrated and desalted using a VanGuard™ CSH C18 pre-column (1.7 μm, 2.1 x 5 mm; Waters Corporation, Miliford, MA), and separated using an ACQUITY UPLC™ CSH C18 column (1.7 μm, 1 x 100 mm, Waters Corporation, Miliford, MA). Labelled peptides were separated over an 8 min gradient of 5-30% acetonitrile at 40 μL/min and 0°C. After each run, the pepsin column was manually cleaned with two consecutive washes of 1.5% formic acid, 5% acetonitrile, 1.5 M guanidinium chloride, pH 1.6. Blank injections were performed after each sample to confirm the absence of peptide carry-over.

The LC flow was directed to a Synapt™ G2-Si HDMS™ mass spectrometer (Waters Corporation) equipped with a standard electrospray ionization source (ESI). Mass accuracy was ensured by continuously infusing a Glu-1-Fibrinogen solution (100 fmol/μL in 50% acetonitrile, 0.1% formic acid) through the reference probe of the ESI source at a flow rate of 3 µL/min. Mass spectra were acquired in positive-ion and resolution mode over the 50–1950 m/z range. CaM peptic peptides were identified in undeuterated samples by a combination of data independent acquisition (MS^E^) and exact mass measurement (below 10 ppm mass error) using the same chromatographic conditions than for the deuterated samples. Four distinct MS^E^ trap collision energy ramps were employed to optimize the efficiency of the fragmentation: 10-30V (low), 15-35V (medium), 20-45V (high), and 10-45V (mixed mode).

**HDX-MS data processing**

DynamX 3.0 (Waters Corporation, Miliford, MA) was used to extract the centroid masses of all peptides selected for local HDX-analyses; only one charge state was considered per peptide. The final peptide map of CaM was refined in DynamX 3.0 using the following Protein Lynx Global Server (PLGS) import options: minimum intensity: 3000; sum of products per amino acid: 0.15; sum of intensity for products: 1000; minimum PLGS score: 6.5; maximum MH+ error (ppm): 5; file threshold: 2. A total of 40 peptides covering 93.2% of the CaM sequence were selected for HDX-MS based on their signal intensity and signal over noise ratio (Figure S3). No adjustment was made for back-exchange; results are therefore reported as relative deuterium exchange levels expressed in either mass unit or fractional exchange. Overlapping CaM peptides were only used to improve the spatial resolution if their back exchange values were identical and ≤ at 10% (Note: the fully deuterated CaM sample was acquired in a previous study using identical experimental conditions). Fractional exchange data was calculated by dividing the experimentally measured uptake by the theoretically maximum number of exchangeable backbone amide hydrogens that could be replaced into each peptide (taking into account the final excess of deuterium present in the labeling mixture). MEMHDX was used to visualize and statistically validate HDX results (Wald test, false discovery rate sets to 5%) ^28^.

**Nuclear Magnetic Resonance**

Samples of ^15^N or ^15^N/^13^C labeled Holo-CaM (Giotto Biotech, Italy) were prepared in 20 mM HEPES pH 7.0, 100 mM NaCl, 2 mM CaCl_2_, supplemented with appropriate amounts of D_2_O/DMSO-d6 (Eurisotop, France). Protein concentration ranged between 30 and 270 µM. For Holo-CaM-CDZ interaction studies, CDZ dissolved in DMSO-d6 was added to Holo-CaM samples. The final DMSO-d6 concentration varied between experiments but was always lower than 7%. We checked by using NMR ^1^H-^15^N correlation spectroscopy that Holo-CaM spectra, and hence the protein structure and dynamics, were not affected by DMSO up to 10% concentration.

NMR experiments were run on a 600 MHz Avance III HD spectrometer (Bruker BioSpin, Billerica, USA) equipped with a triple resonance (^1^H/^13^C/^15^N) cryogenically cooled probe. Spectra were recorded with Topspin 3.6.3 (Bruker), processed with NMRPipe ^29^ and analyzed with CCPNMR analysis 2.4.3 ^30^. Experiments were performed at 37°C and referenced to the sodium salt of 4,4-dimethyl-4-silapentane-1-sulfonic acid.

Backbone (^1^H,^15^N, CA) and CB chemical shifts of the free (260 µM) and CDZ bound forms (265 µM) of Holo-CaM were assigned using standard two- and three-dimensional experiments: ^13^C and ^15^N HSQC (heteronuclear single quantum coherence) ^31^, ^1^H-^15^N SOFAST-HMQC, and the BEST (band-selective excitation short transient) versions of the HNCACB, CBCA(CO)NH, HNCA and HN(CO)CA pulse sequences implemented in NMRLib 2.0 ^32^. While for free Holo-CaM, all residues could be assigned, for the Holo-CaM:CDZ complex there were 11 missing resonances in ^1^H-^15^N correlation spectra and 22 were shifted and typically too broad to permit assignment.

Secondary structures of the free and bound forms were estimated from backbone ^1^H,^15^N, CA and side chain CB chemical shifts using Talos-N ^33^. The secondary structures of Holo-CaM thus obtained, were compared to the secondary structures of Holo-CaM (^34^, PDB codes 1J7O and 1J7P for the N- and C-lobes, respectively) and Holo-CaM:CDZ complexes determined in this work as established by DSSP^35^.

Backbone and CB chemical shifts were also used to calculate the order parameter (S^2^) of isolated Holo-CaM to analyze the amplitude of motions on the ns-ps time range. S^2^ values were determined using the RCI software^36^.

The chemical shift perturbation (CSP) upon CDZ binding to Holo-CaM was determined from the differences of ^15^N (∆δ_N_) and ^1^HN (∆δH_N_) chemical shifts of the free and bound forms using the formula:

$$CSP = \sqrt{{{(0.159 \times{\Delta\delta}_{N})}^{2} + \Delta\delta}_{HN}^{2}}$$

The translation (d) and rotational diffusion (τ_c_) of Holo-CaM and the effect of CDZ binding at different Holo-CaM:CDZ ratios were evaluated on ^15^N labeled Holo-CaM samples (± CDZ) by ^1^H detected 1D ^15^N-edited ^1^H pulsed-field gradient self-diffusion experiments coupled to the ^1^H-^15^N SOFAST-HMQC sequence, and by ^15^N cross-correlated spin relaxation experiments (TRACT, TROSY for rotational correlation times ^37^) as implemented in NMRLib 2.0. To take into account the effect on viscosity of different D_2_O and DMSO concentrations, results were extrapolated to 100% H_2_O using the viscosities of the corresponding H_2_O, D_2_O and DMSO mixtures.

The internal dynamics of Holo-CaM were analyzed by determining the ^15^N longitudinal (T_1_) and transverse (T_2_) relaxation rates and the steady state ^1^H-^15^N heteronuclear nOe (nuclear Overhauser enhancement) relaxation parameters from 2D ^1^H-^15^N experiments ^38^. The concentration of Holo-CaM was 260 µM (5% D_2_O) for the free protein, 265 µM for the 1:1 and 65 µM for the 1:3 Holo-CaM:CDZ samples (4% DMSO). Series of 5-6 spectra with a 5 s recovery delay between scans and different relaxation delays were recorded for T_1_ and T_2_ measurements. The ^1^H-^15^N nOe experiments were run with a recovery delay of 3 s and a saturation/no saturation delay of 4 s. Measurement errors were estimated from the spectra noise standard deviation. Fits to exponential decays and Monte Carlo evaluation of fit errors and confidence levels were performed with in-house Python scripts.

**Statistical Analysis of the experimental datasets**

2D-SAXS data were radially averaged, normalized to the intensity of the incident beam and put on an absolute scale using the scattering from water ^39^ before buffer scattering subtraction. All these operations were performed using the programs FoxTrot (courtesy of SWING beamline) and Primus (<https://www.embl-hamburg.de/biosaxs/primus.html>) ^40^. The resulting 1D scalar scattering intensity profiles are represented by the normalized intensities and their associated standard deviations (SD). Identical frames under the main elution peak were selected using Cormap ^41^ and averaged for further analysis. The agreement between experimental data and calculated intensities from models was evaluated using the reduced χ^2^ metric^40^.

X-ray data collection, processing and model refinement statistics corresponding to the X-ray structures are summarized in Table S2. The software packages used are autoPROC, Phaser, Coot, Buster and Pymol, as described in the Crystallography section.

The statistical analysis for the HDX-MS experiments is described below. A summary of the HDX-MS experiments is provided in Table S4 and in the HDX-MS analysis section. Pre-processing of data: one unique charge state was considered per peptide (user selection). The quality determination of the dataset was accomplished by MEMHDX (<http://memhdx.c3bi.pasteur.fr>) to evaluate the agreement across replicates. Data presentation: Logit representation is used for Figure S11.

For Figure S11, each dot reported on the uptake differential plots corresponds to the average value of three independent replicates. The repeatability of the measurement determined for each state (pooled standard deviation) using the 46 selected peptides and the 966 unique MS data points per state is reported in Table S4. Sample size (n) for each statistical analysis: Triplicate labeling experiments were performed for each time point (7 time points including unlabeled controls) and condition for all HDX-MS analyses (independent technical replicates). Considering 46 peptides, 1 charge state, 3 replicates, 7 time points and 2 conditions, the complete HDX-MS datasets contains n= 1932 unique data points. Statistical methods used to assess significant differences with sufficient details: To consider the time dependency of the exchange reaction, two distinct p-values were calculated per peptide using two individual Wald tests. The FDR was set to 5% (p < 0.05). Software used for statistical analysis: MEMHDX software (<http://memhdx.c3bi.pasteur.fr>).

The statistical analysis for NMR experiments was performed as follows. Measurement errors were estimated from the spectra noise standard deviation. Fits to exponential decays and Monte Carlo evaluation of fit errors and confidence levels were performed with in-house Python scripts. Error bars were obtained from 1000 Monte Carlo simulations considering one noise standard deviation.

**Supplementary Figures**

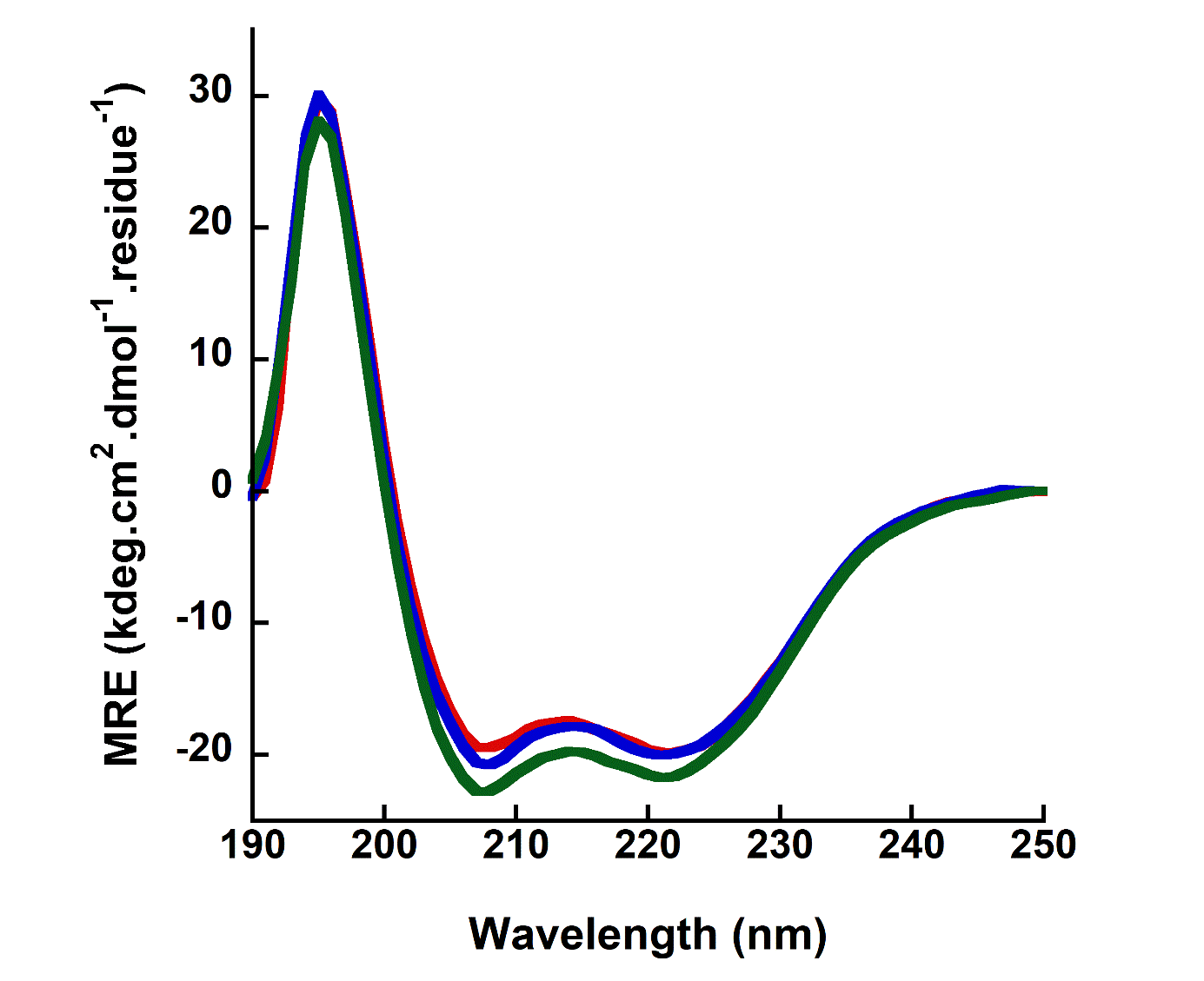

**Supplementary Figure S1**: **Synchrotron radiation far-UV circular dichroism.** Holo-CaM (60 µM) in buffer B with 0.91% final DMSO in the absence (red) and in the presence of 66 µM of CDZ (blue, Holo-CaM:CDZ molar ratio of 1:1.1) or 132 µM of CDZ (green, Holo-CaM:CDZ molar ratio of 1:2.2).

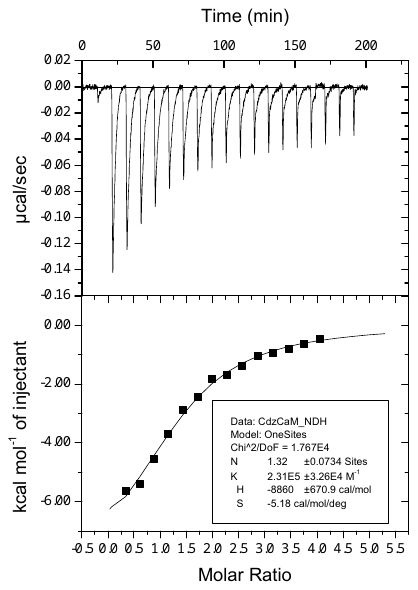

**Supplementary Figure S2**: **Isothermal titration calorimetry of Holo-CaM:CDZ.** A solution of 8 µM Holo-CaM was loaded in the reaction cell. A solution of 200 µM CDZ was loaded into the syringe and then injected into the reaction cell. Heats of dilutions were measured by injecting 200 µM of CDZ into buffer B and were subtracted from the heat of reaction of the Holo-CaM titration by CDZ. The titration profiles were analyzed using the Origin7.0 software (OriginLab) to determine the thermodynamic parameters. In the example shown in this figure, the dissociation constant, K_D_, is 4.3 µM and the stoichiometry is 1.3. The average values of K_D_ and stoichiometry (n=6 experiments) are 3 ± 2 µM and 1.2 ± 0.5, respectively.

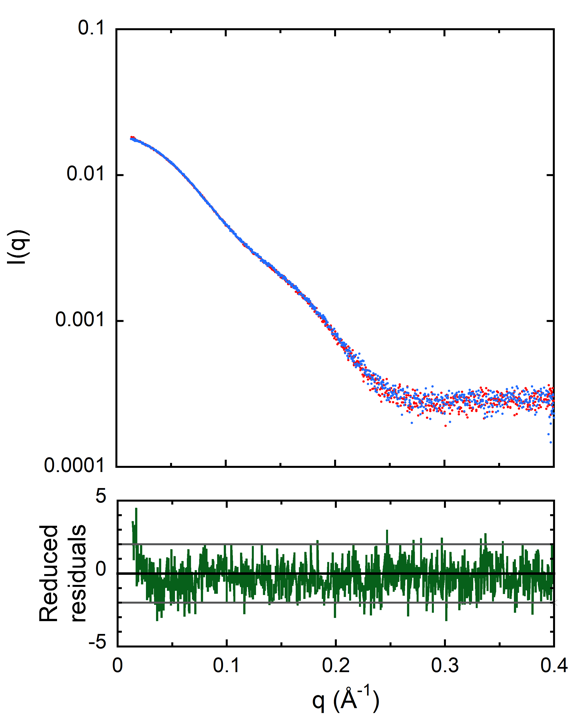

**Supplementary Figure S3:** **Analysis of DMSO effect on Holo-CaM using SAXS.** Scattering patterns of Holo-CaM in buffer B in the absence (red) and in the presence of 5% DMSO (blue) together with the reduced residuals distribution (green line).

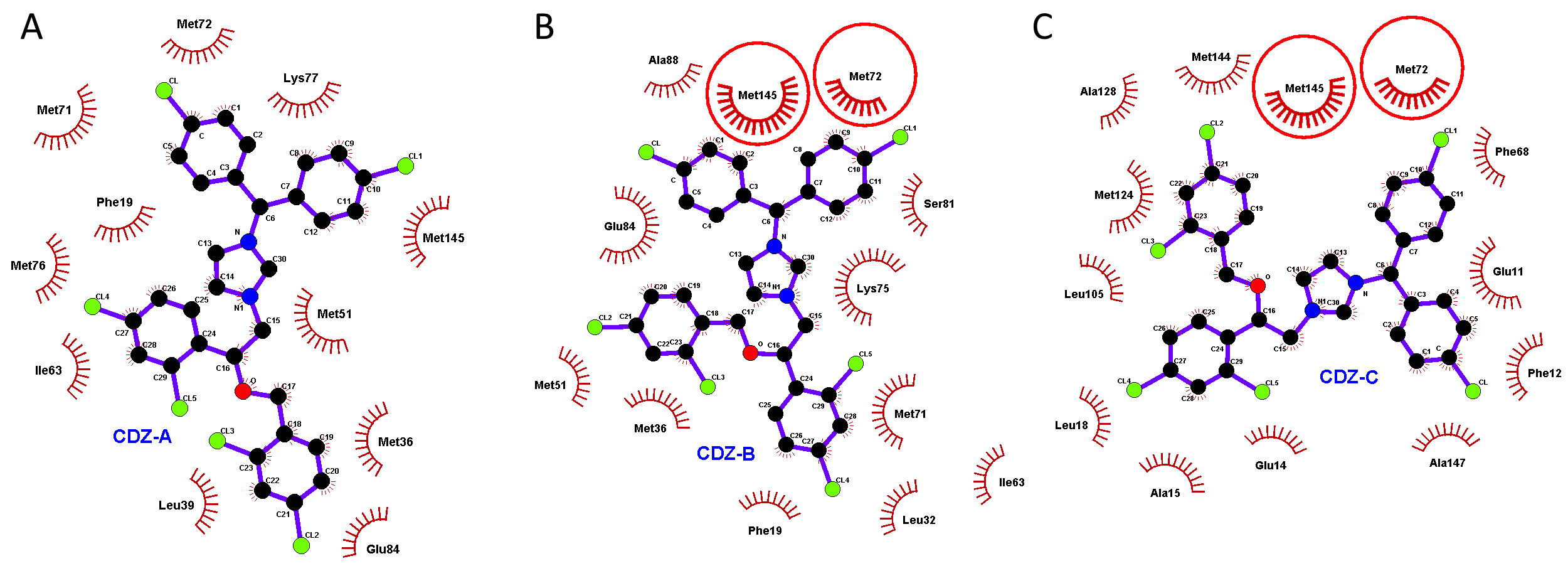

**Supplementary Figure S4**: **Interactions between CDZ and Holo-CaM residues visualized using Ligplot+**. **A**. Ligplot of CDZ from the crystallographic structure of the 1:1 Holo-CaM:CDZ complex (pdb access code 7PSZ) and its interacting residues. Most of these residues are located in the C-terminal part of N-CaM and in the vicinity of the central linker (residues 77-81). **B. and C.** Ligplot of the two CDZ molecules from the Holo-CaM:CDZ 1:2 crystallographic structure (pdb access code 7PU9). Whereas CDZ-B interacts with almost the same residues than CDZ-A mainly located in the N-terminal lobe, CDZ-C interacts with residues in both lobes. Methionine residues 72 and 145 (circled in red) interact with both CDZ molecules, as well as with CDZ-A in the 1:1 complex. The vast majority of the Holo-CaM:CDZ interactions are hydrophobic.

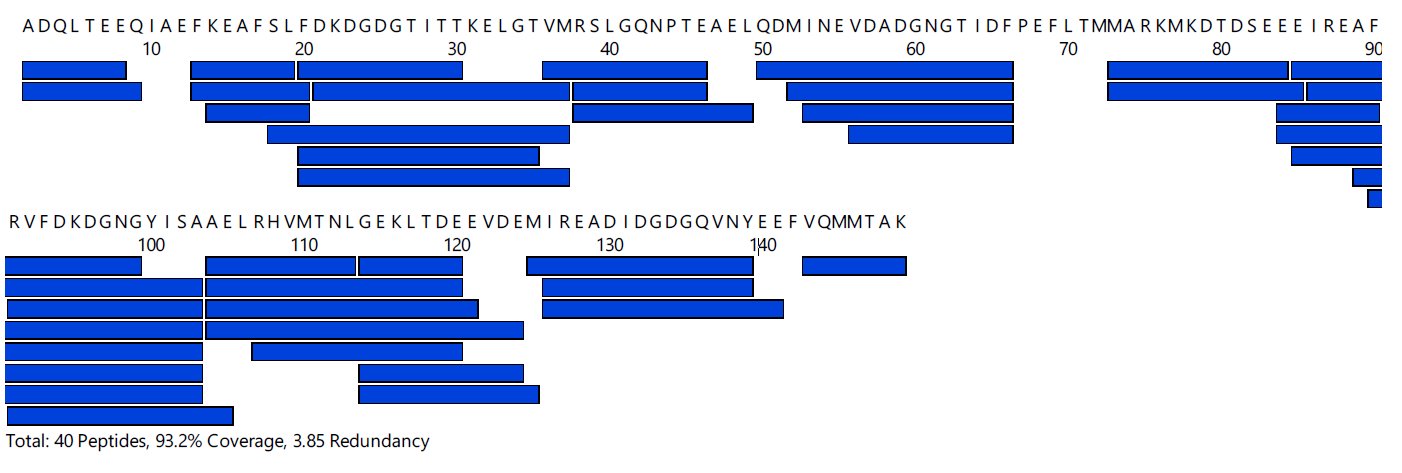

**Supplementary Figure S5**: Peptide map of CaM generated after 2 min pepsin digestion at 20°C and pH 2.5. Each blue bar corresponds to a unique peptide. A total of 40 peptides covering 93.2% of the protein sequence with a 3.85 redundancy value were selected for HDX-MS.

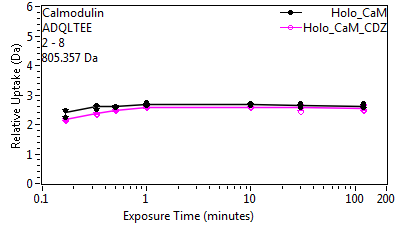

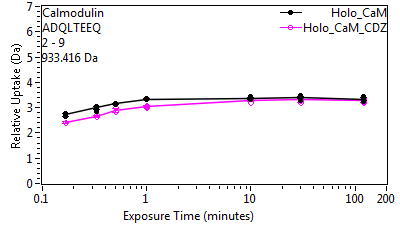

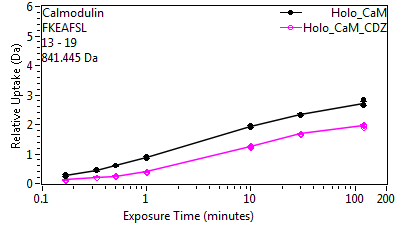

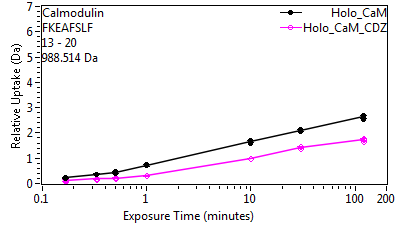

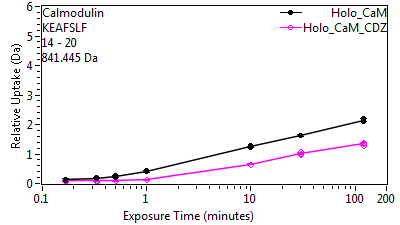

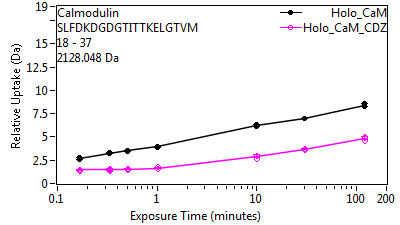

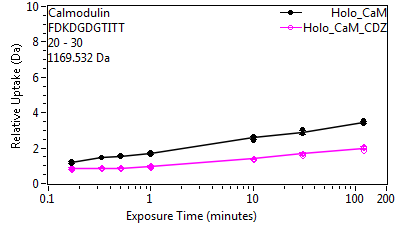

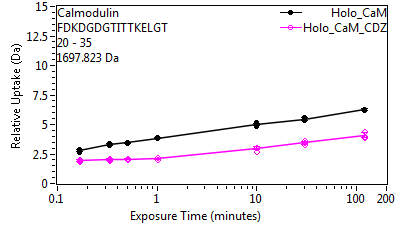

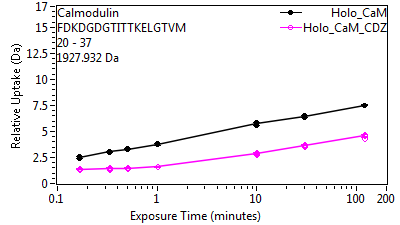

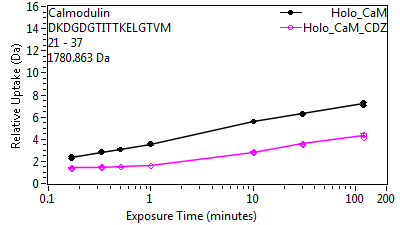

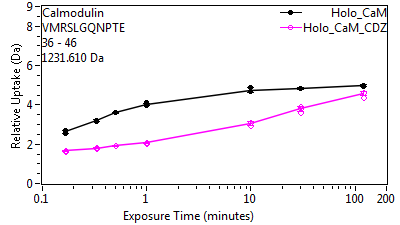

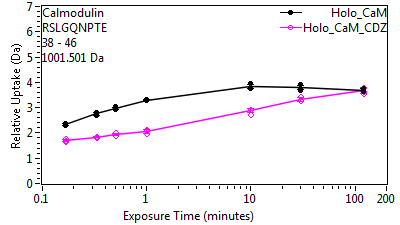

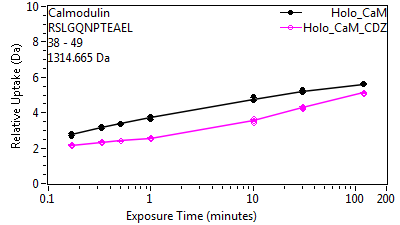

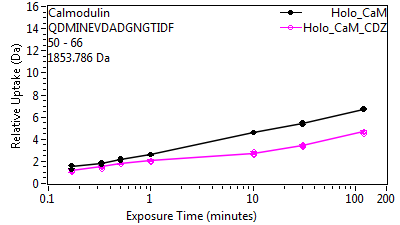

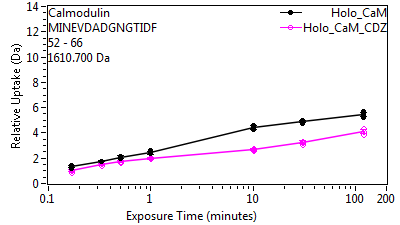

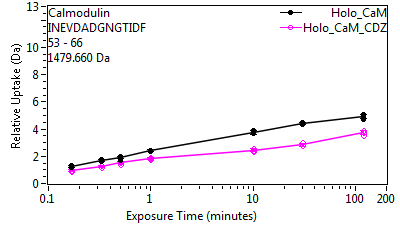

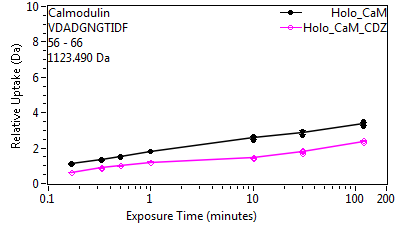

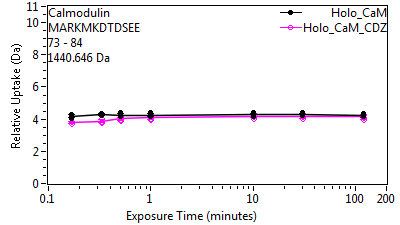

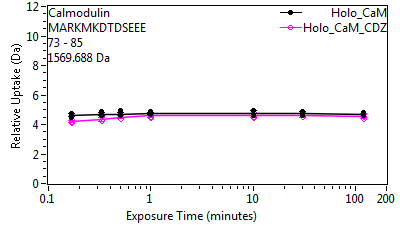

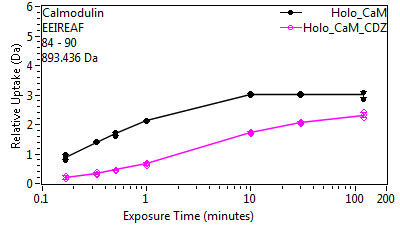

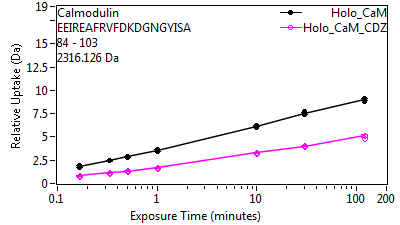

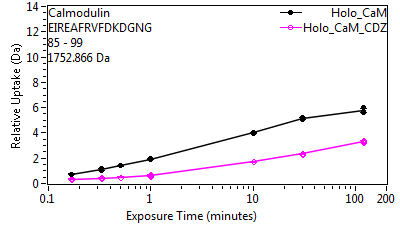

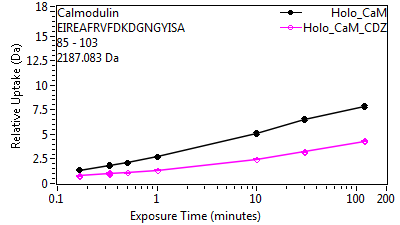

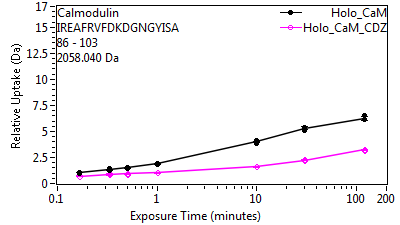

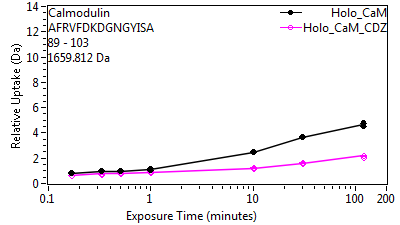

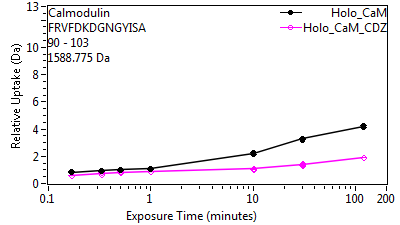

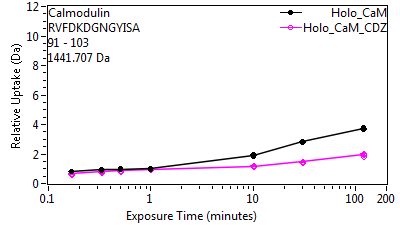

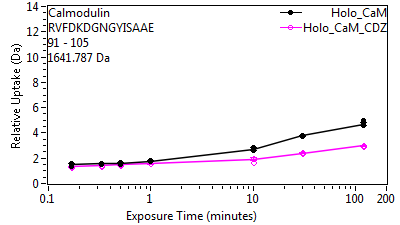

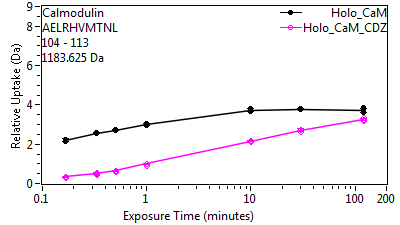

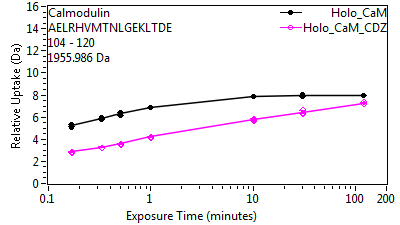

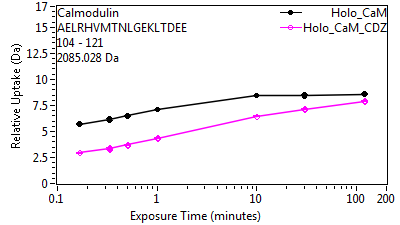

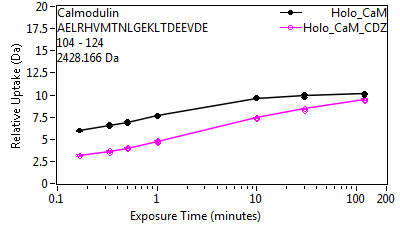

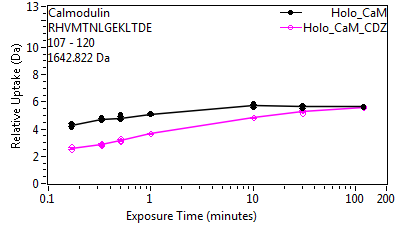

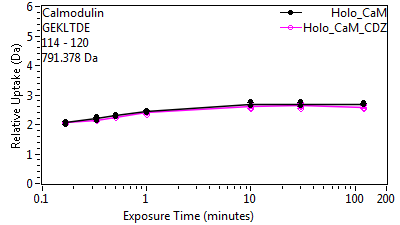

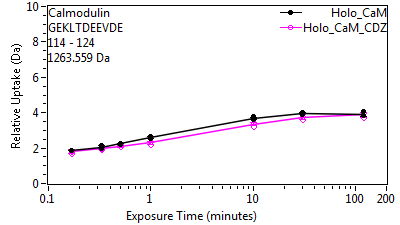

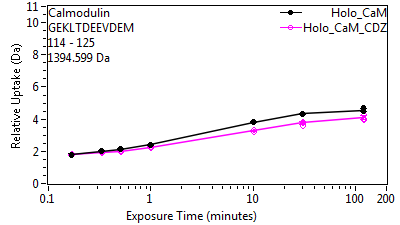

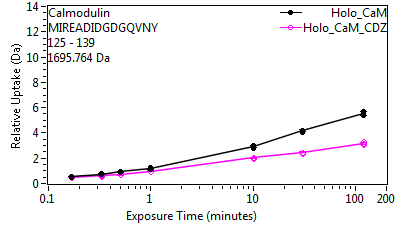

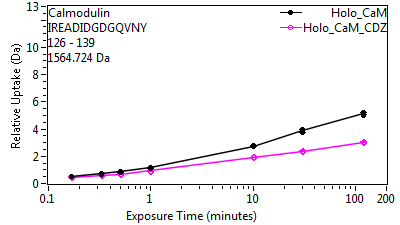

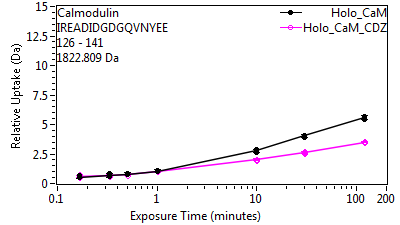

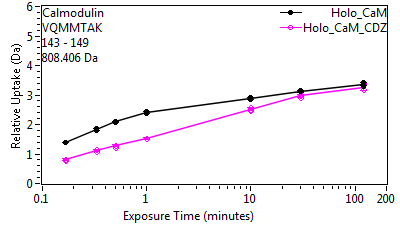

**Supplementary Figure S6:** Deuterium uptake plots for each individual CaM peptides across all conditions.

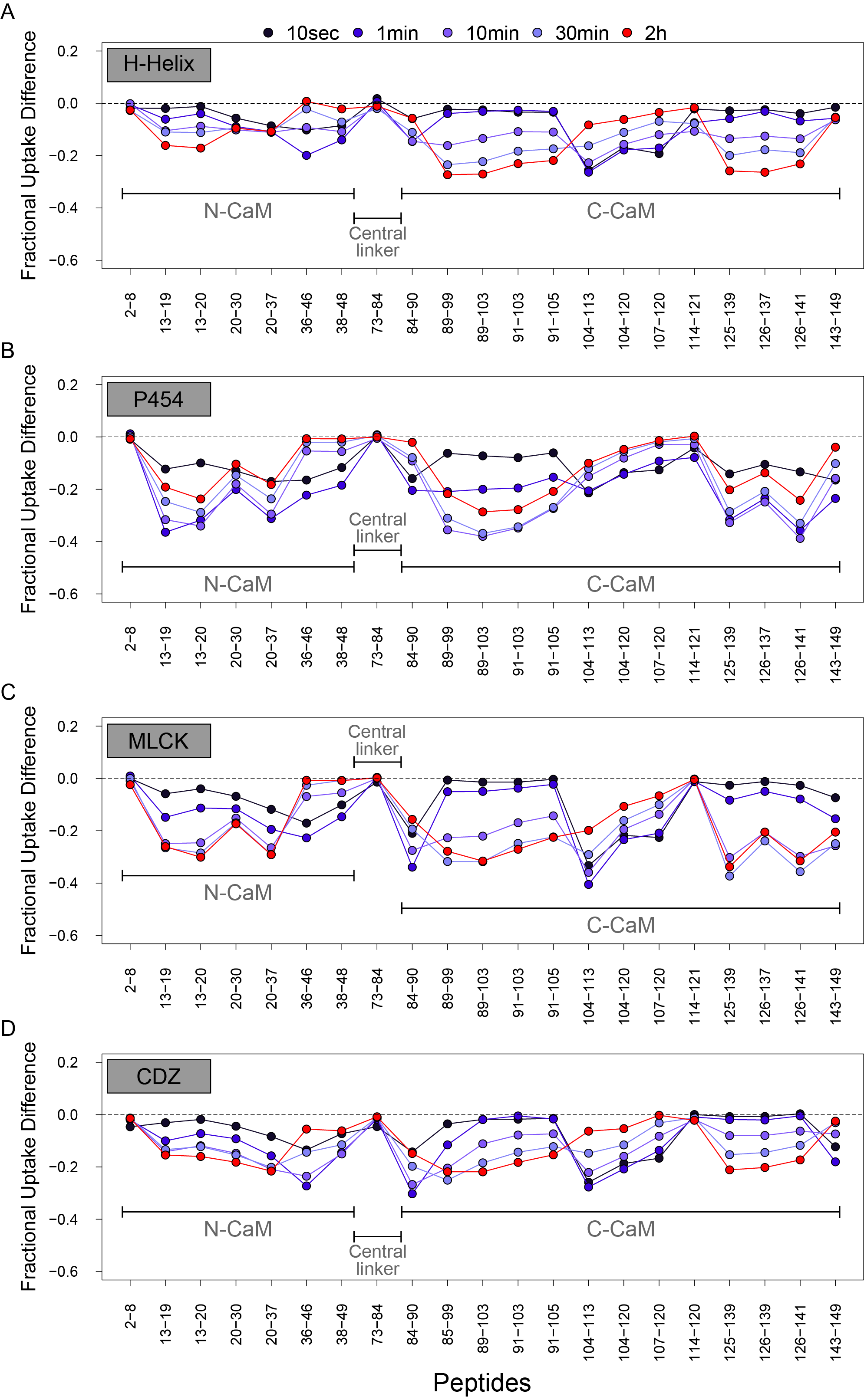

**Supplementary Figure S7**: **Comparison of the differential HDX patterns within Holo-CaM upon H-helix (A), P454 (B), MLCK (C) and CDZ (D) binding.** Each differential fractional uptake plot shows the differences in uptake calculated between the CDZ-bound and the free Holo-CaM states at each time point, and for each selected peptide. A negative uptake difference value indicates a ligand-induced protective effect (i.e., reduction in solvent accessibility).

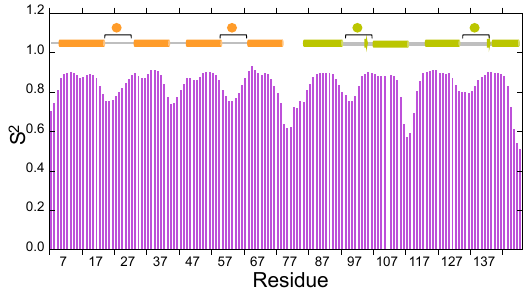

**Supplementary Figure S8. Order parameter (S^2^) reflecting the amplitude of motions on the ns-ps time scales of free Holo-CaM.** The secondary structure (helix=cylinder, strand=arrow) and calcium ion loops (spheres and square brackets) are schematized on the top of each graph, with the N- and C-lobes colored in orange and green, respectively. Backbone amide S^2^ values were calculated from backbone chemical shifts using RCI. The order parameter, which varies from 0 to 1, reflects the amplitude of internal motions of the backbone NH bonds on the ns-ps time scale (0: unrestricted motions; 1: no internal motions). The N and C-termini, calcium binding loops, the loop between helices H6 and H7 and the interlobe linker region (highlighted in grey) show fast high amplitude motions.

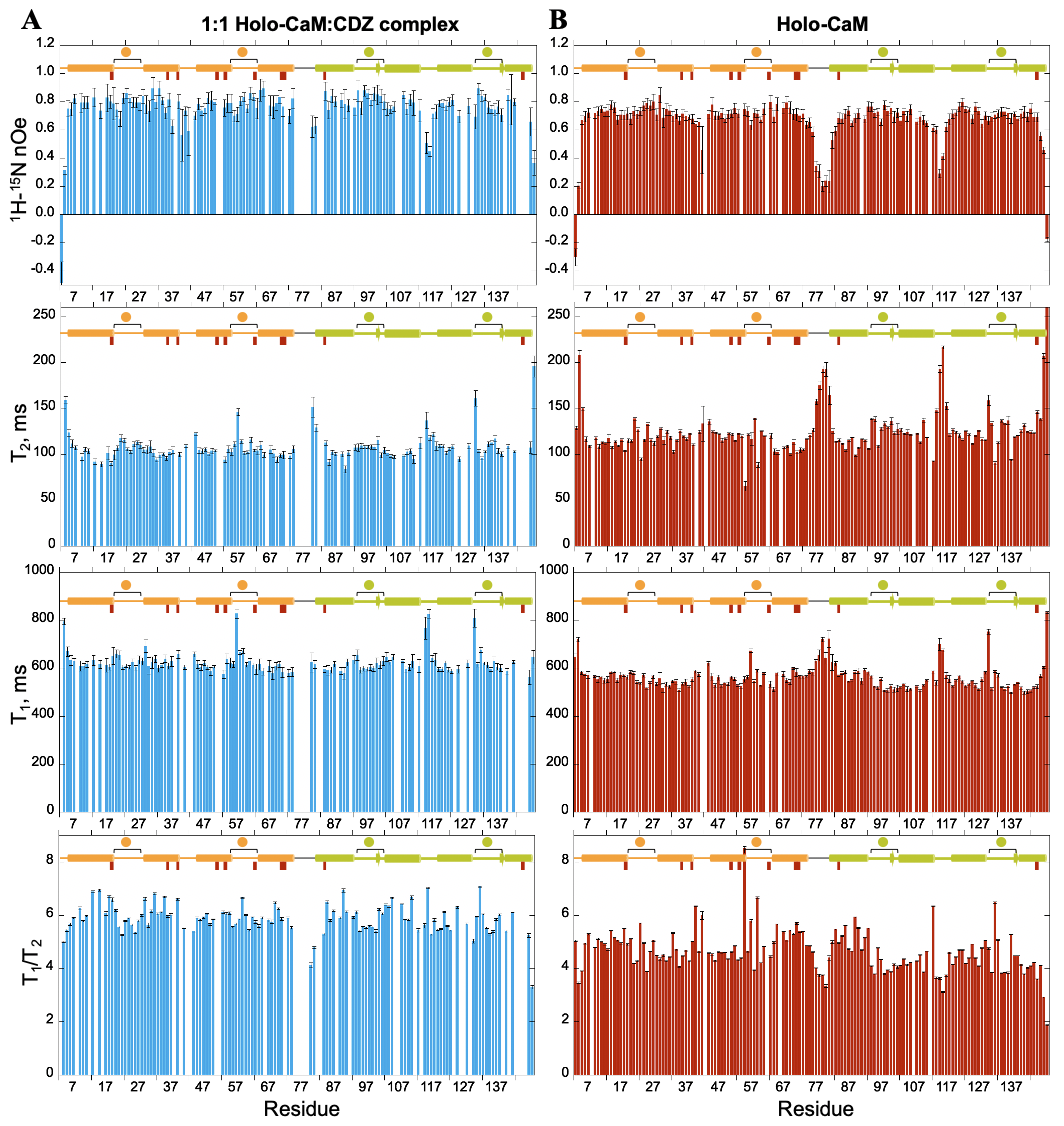
**Supplementary Figure S9**: **Holo-CaM internal dynamics from ^15^N relaxation data.** T_1_, T_2_ T_1_/T_2_ and ^1^H-^15^N nOe of Holo-CaM isolated **A.** or in a 1:1 complex with CDZ **B.** as a function of the residue number. Data were collected at 37 °C on a 600 MHz spectrometer. The secondary structure (helix=cylinder, strand=arrow) and calcium ion loops (spheres and square brackets) are schematized on the top of each graph, with the N- and C-lobes colored in orange and green, respectively. Lacking data are due to either proline residues, signal overlap and unassigned (shifted very weak or absent exchanged-broadened signals). The positions of contacting residues in the X-Ray 1:1 Holo-CaM:CDZ complex structure, as defined by Ligplot+ are represented by wine rectangles. In Holo-CaM, the relaxation parameters (high T_2_, low T_1_, low T_1_/T_2_ and low ^1^H-^15^N nOes) of residues in the linker region (77-81) and flanking residues in α-helices H4 and H6 are characteristic of high amplitude motions on the ns-ps time scale, with an increased flexibility around its mid-point (79T-80D). In contrast, the relaxation parameters of the α-helical elements of the N- and C-lobes are indicative of ordered regions in a globular protein. The N- (residues 3-4) and C-terminal (146-148) residues are disordered and loops between helices H2-H3 (39-42) and 6-7 (110-118) show increased mobility with respect to the other residues in both lobes’ calcium binding EF-hands, as assessed by high T_2_ and low ^1^H-^15^N nOe values. Finally, residues 55V and 59G in Ca^2+^ binding loop 2, as well as residues 132G and 137N in Ca^2+^ binding loop 4, display low T_2_ values revealing conformational exchange phenomena on the ms-µs time scale. Our data agree very well with published data on Holo-CaM obtained at 35 °C and 500 MHz ^42^), in which the first demonstration that the N- and C-lobes move independently of each other was done. Error bars were obtained from 1000 Monte Carlo simulations considering one noise standard deviation.

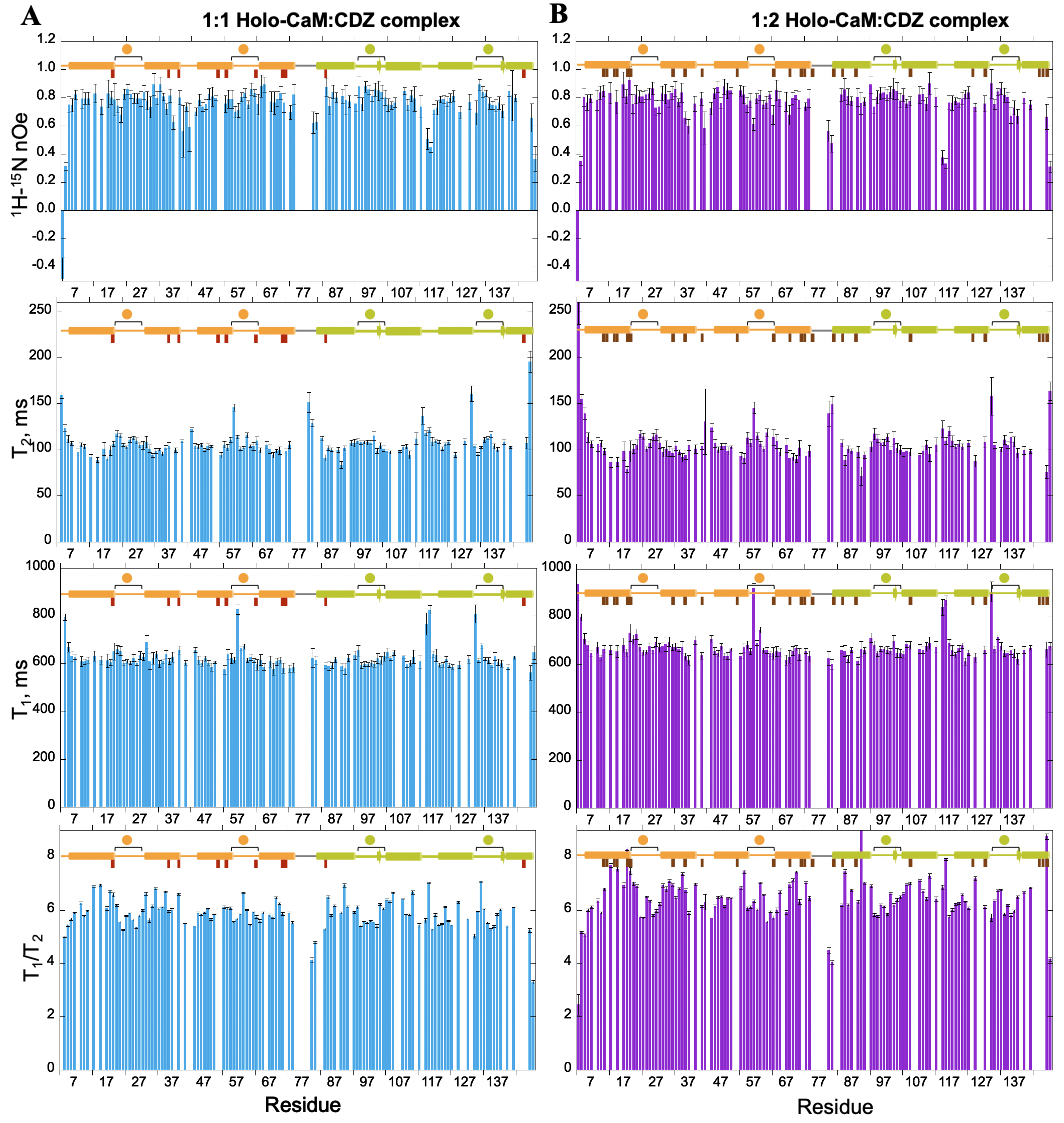
**Supplementary Figure S10**: **Comparison of Holo-CaM internal dynamics from ^15^N relaxation data in complex with CDZ.** T_1_, T_2_ T_1_/T_2_ and ^1^H-^15^N nOe of Holo-CaM in complex with 1 **A.** or 2 **B.** CDZ molecules as a function of the residue number. The T_1_/T_2_ ratios, which in the absence of contributions from conformational exchange and for residues in rigid structures (nOe ≥ 0.65), are directly related to the tumbling correlation time of a molecule. Data were collected at 37 °C on a 600 MHz spectrometer. Data for the 1:2 complex was obtained with a 3-fold excess of CDZ relative to Holo-CaM. The secondary structure (helix=cylinder, strand=arrow) and calcium ion loops (spheres and square brackets) are schematized on the top of each graph, with the N- and C-lobes colored in orange and green, respectively. Lacking data are due to either proline residues, signal overlap and unassigned (very weak shifted or absent exchanged-broadened signals). The positions of contacting residues in the X-Ray 1:2 Holo-CaM:CDZ complex structure, as defined by Ligplot+ are represented by wine (1:1 complex) and maroon (1:2 complex) rectangles. Error bars were obtained from 1000 Monte Carlo simulations considering one noise standard deviation. The nOe, T_1_ and T_2_ profiles are very similar in 1 or 2 CDZ-loaded Holo-CaM, albeit some residues showing shorter/longer T_2_ values indicative of different conformational exchange rates and slightly higher T_1_ values for the 1:3 sample, most likely due to the higher molecular weight of the complex.

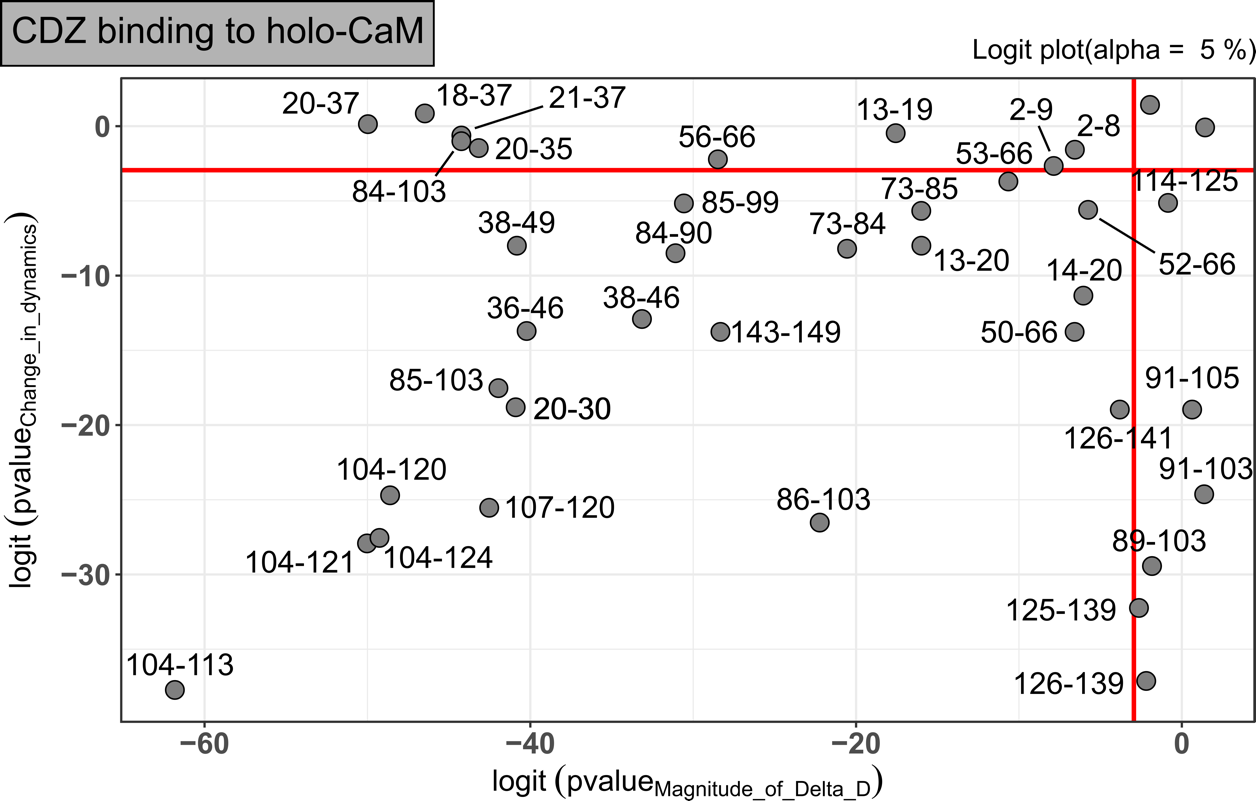

**Supplementary Figure S11**: **Statistical analysis performed with MEMHDX**. Logit plot showing the statistical results obtained with all individual CaM peptides in the presence of a 32x molar excess CDZ. The two red lines correspond to the statistically significant threshold used during the statistical analysis (FDR sets to 5%, Wald test, biological threshold sets to 2%). Peptides showing no statistically significant uptake differences between states are displayed on the top right-hand rectangle created by the two red lines. Statistically significant peptides are labelled.

| **Data collection parameters** | | | |
| --- | --- | --- | --- |
| Instrument | Beamline SWING (Synchrotron SOLEIL) | | |
| Detector | EigerX4M in vacuum (Dectris) | | |
| Sample to detector distance | 2.00 m | | |
| Beam geometry | 400 µm x 200 µm | | |
| Wavelength (Å) | 1.033 | | |
| q-range (Å^-1^) | 0.0041 < q < 0.50 | | |
| Exposure time (ms) / reading time (ms) | 990 / 10 | | |
| Temperature (K) | 288 | | |
| Sample | Holo-CaM | Holo-CaM (prepared in 5% DMSO) | Holo-CaM:CDZ (prepared in 7% DMSO) |
| Molar extinction coefficient  (M^-1^.cm^-1^) | 2 980 | | |
| CaM molecular mass^(1)^ (Da) | 16 706 | | |
| Initial CaM concentration (µM) | 500 | 500 | 333 |
| Initial CDZ concentration (µM) | 0 | 0 | 1050 |
| Running buffer of the SEC | Buffer B | | |
| **Structural parameters** | | | |
| I(0) Guinier (cm^-1^) | 0.0182 | 0.0203 | 0.0129 |
| R_g_ Guinier (Å) | 21.98 ± 0.06 | 21.83 ± 0.03 | 16.89 ± 0.04 |
| I(0) P(r) (cm^-1^) | 0.183 | 0.0204 | 0.0129 |
| R_g_ P(r) (Å) | 22.49 ± 0.03 | 22.39 ± 0.04 | 16.83 ± 0.03 |
| D_max_ (Å) | 74 | 72 | 52 |
| **Data reduction and analysis software** | | | |
| FOXTROT, ATSAS, RAW, US-SOMO, GNOM | | | |
| **Modeling software** | | | |
| DENSS^14^, EOM ^15,16^Tria, G., Mertens, H. D. T., Kachala, M. & Svergun, D. I. (2015) Advanced ensemble modelling of flexible macromolecules using X-ray solution scattering. *IUCrJ* 2, 207-217. | | | |

**Supplementary Table S1: SAXS data collection and scattering derived parameters.**

(1) Molecular mass of calmodulin in the absence of the N-terminal Methionine residue. The expected averaged molecular mass from Protpi server (<https://www.protpi.ch/Calculator/ProteinTool>) is 16 706.25 Da. The averaged molecular mass measured by mass spectrometry is 16 706.18 +/-0.07 Da (delta = 0.07 Da, *i.e.*, 4.2 ppm).

| **Complexes** | Holo-CaM:CDZ  (molar ratio of 1:2.2) | Holo-CaM:CDZ  (molar ratio of 1:10) |
| --- | --- | --- |
| **Pdb code** | 7PSZ | 7PU9 |
| **Beamline** | PROXIMA-2A | PROXIMA-1 |
| **Wavelength** | 0.9801 | 0.8266 |
| **Resolution range** | 30.91 - 1.898 (1.966 - 1.898) | 34.08 - 2.279 (2.36 - 2.279) |
| **Space group** | C 1 2 1 | P 6_1_ 2 2 |
| **PDB access code** | 7PSZ | 7PU9 |
| **Unit cell** | 66.848 36.127 68.382  90 116.738 90 | 39.351 39.351 336.92  90 90 120 |
| **Total reflections** | 78268 (7850) | 278336 (29794) |
| **Unique reflections** | 11450 (1097) | 8006 (751) |
| **Multiplicity** | 6.8 (7.2) | 34.8 (39.7) |
| **Completeness (%)** | 97.55 (96.39) | 99.93 (99.87) |
| **Mean I/sigma(I)** | 17.21 (2.57) | 17.84 (2.71) |
| **Wilson B-factor** | 37.33 | 52.14 |
| **R-merge** | 0.06148 (0.8184) | 0.1477 (1.307) |
| **R-meas** | 0.06686 (0.8823) | 0.15 (1.324) |
| **R-pim** | 0.02585 (0.3274) | 0.02589 (0.2087) |
| **CC1/2** | 0.998 (0.906) | 0.999 (0.909) |
| **CC*** | 1 (0.975) | 1 (0.976) |
| **Reflections used in refinement** | 11450 (1095) | 8006 (750) |
| **Reflections used for R-free** | 583 (45) | 406 (25) |
| **R-work** | 0.2281 (0.3077) | 0.2341 (0.2600) |
| **R-free** | 0.2574 (0.3430) | 0.2663 (0.2775) |
| **CC(work)** | 0.947 (0.802) | 0.904 (0.862) |
| **CC(free)** | 0.946 (0.442) | 0.932 (0.902) |
| **Number of non-hydrogen atoms** | 1281 | 1259 |
| **macromolecules** | 1149 | 1134 |
| **ligands** | 89 | 84 |
| **solvent** | 43 | 41 |
| **Protein residues** | 145 | 144 |
| **RMS(bonds)** | 0.011 | 0.014 |
| **RMS(angles)** | 1.45 | 1.48 |
| **Ramachandran favored (%)** | 99.30 | 97.89 |
| **Ramachandran allowed (%)** | 0.70 | 2.11 |
| **Ramachandran outliers (%)** | 0.00 | 0.00 |
| **Rotamer outliers (%)** | 1.59 | 6.50 |
| **Clashscore** | 7.35 | 3.51 |
| **Average B-factor** | 45.22 | 60.49 |
| **macromolecules** | 43.24 | 60.53 |
| **ligands** | 69.12 | 59.36 |
| **solvent** | 48.66 | 61.64 |

**Supplementary Table S2: Crystallographic data collection and refinement statistics.**

Statistics for the highest-resolution shell are shown in parentheses.

|  |  | **Holo-CaM:CDZ 1:1** | | **Holo-CaM:CDZ 1:2** | | | |  | **Effect in HDX-MS** |
| --- | --- | --- | --- | --- | --- | --- | --- | --- | --- |
|  |  | **CDZ A** | | **CDZ B** | | **CDZ C** | |  |  |
|  | **N-Lobe** |  | |  | | 11 | Glu |  |  |
|  |  |  |  |  |  | 12 | Phe |  |  |
|  |  |  |  |  |  | 14 | Glu |  |  |
|  |  |  |  |  |  | 15 | Ala |  |  |
|  |  |  |  |  |  | 18 | Leu |  |  |
|  |  | 19 | Phe | 19 | Phe |  | |  |  |
|  |  |  | | 32 | Leu |  |  |  |  |
|  |  | 36 | Met | 36 | Met |  |  |  |  |
|  |  | 39 | Leu |  | |  |  |  |  |
|  |  | 51 | Met | 51 | Met |  |  |  |  |
|  |  | 63 | Ile | 63 | Ile |  |  |  |  |
|  |  |  | |  | | 68 | Phe |  |  |
|  |  | 71 | Met | 71 | Met |  | |  |  |
|  |  | 72 | Met | 72 | Met | 72 | Met |  |  |
|  | **Linker region** |  |  | 75 | Lys |  | |  |  |
|  |  | 76 | Met |  | |  |  |  |  |
|  |  | 77 | Lys |  |  |  |  |  |  |
|  |  |  |  | 81 | Ser |  |  |  |  |
|  | **C-Lobe** | 84 | Glu | 84 | Glu |  |  |  |  |
|  |  |  | | 88 | Ala |  |  |  |  |
|  |  |  |  |  | | 105 | Leu |  |  |
|  |  |  |  |  |  | 124 | Met |  |  |
|  |  |  |  |  |  | 128 | Ala |  |  |
|  |  |  |  |  |  | 144 | Met |  |  |
|  |  | 145 | Met | 145 | Met | 145 | Met |  |  |
|  |  |  |  |  |  | 147 | Ala |  |  |

**Supplementary Table S3**: Detailed location of residues interacting with CDZ according to Ligplot+ and comparison with HDX-MS data. CDZ-A is the CDZ molecule from the crystallographic structure of the 1:1 Holo-CaM:CDZ complex (pdb 7PSZ, Holo-CaM:CDZ_A_). CDZ-B and CDZ-C are the CDZ molecules from the crystallographic structure of the 1:2 Holo-CaM:CDZ complex (pdb 7PU9, Holo-CaM:CDZ_BC_). The effects observed by HDX-MS correspond to the average [Δdeuteration] taken from Figures 3 and S6, the scale is from 0 % (blue) to 7.5% (white) to 15 % (red). Grey means no HDX-MS data were obtained.

(1)	The interlobe linker corresponds to residues [76-81] and the linker region to residues [74-82].

| **DATASETS** | **Holo-CaM** | **Holo-CaM:CDZ** |
| --- | --- | --- |
| HDX reaction details   - *pD:* - *T°C:* - *Deuterium level* - *during labelling* - *after quench* - *Calcium:* - *DMSO:* - *CDZ molar excess ^(1)^:* - *% complex during labelling ^(2)^:* | 7.4  RT  80%  26.7%  2 mM  2%  /  / | 7.4  RT  80%  26.7%  2 mM  2%  32x  90 |
| HDX time course analyzed (min) | 0.16, 0.33, 0.5, 1, 10, 30, and 120 min | |
| Number of peptides  Sequence coverage  Average peptide length  Redundancy | 40  93.2%  13.3  3.85 | 40  93.2%  13.3  3.85 |
| Average peptide length / Redundancy ratio | 3.45 | 3.45 |
| Replicates ^(3)^ | 3 | 3 |
| Repeatability (pooled SD, Da) ^(4)^ | 0.06 | 0.06 |
| Significance difference between states ^(5)^ | Wald test, p<0.05; biological threshold set to 3% | |

**Supplementary Table S4**: **HDX-MS data summary**.

(1) Molar excess used in labelling
(2) K_D_ = 3 µM; binding stoichiometry Holo-CaM:CDZ =1:1.2
(3) Independent technical replicates
(4) One unique charge state is used per peptide
(5) MEMHDX software (<https://memhdx.c3bi.pasteur.fr/>)

| **Holo-CaM:CDZ ratio** | **τ_c_ (ns) ^(1)^** |
| --- | --- |
| 1:0.0 | 4.5 ± 0.2 |
| 1:0.5 | 4.7 ± 0.2 |
| 1:1.0 | 5.2 ± 0.3 |
| 1:2.0 | 5.4 ± 0.4 |
| 1:2.7 | 5.7 ± 0.1 |

**Supplementary Table S5**: Rotational correlation time (τ_c_) of Holo-CaM at different Holo-CaM:CDZ ratios. For a protein of a given molecular weight, τ_c_ is expected to be longer for compact conformations relative to compact domains joined by a flexible linker as is the case of Holo-CaM. The τ_c_ value increases at a 1:1 Holo-CaM:CDZ ratio (5.2 ± 0.3 ns) relative to Holo-CaM (4.5 ± 0.2 ns) and is effectively the same at a 1:2 ratio (5.4 ± 0.4 ns). Of note, at 0.5 CDZ equivalents, the τ_c_ (4.7 ± 0.2 ns) corresponds within experimental error to the mean of the free and CDZ-bound protein tumbling times (4.85 ± 0.5 ns), suggesting that free and CDZ-bound (1:1 ratio) forms of Holo-CaM coexist at equivalent concentrations.

(1)	τ_c_ values (± errors) were obtained as described in the Materials and Methods section using the TRACT sequence, which gives tumbling correlation times that are not affected by possible contributions of chemical exchange (between the bound and free forms of a protein or internal conformational exchange) and chemical shift anisotropy.

|  | **Holo-CaM** | | **Holo-CaM : CDZ (1:1)** | | **Holo-CaM : CDZ (1:2)** | |
| --- | --- | --- | --- | --- | --- | --- |
|  | **NMR  Structure** | **NMR Shifts** | **X-Ray** | **NMR Shifts** | **X-Ray** | **NMR Shifts** |
| **Residues in Helix** | **82** | **84** | **85** | **86** | **85** | **86** |
| **Residues in Strands** | **4** | **10** | **4** | **12** | **8** | **12** |
| **Helix (%)** | **55** | **57** | **57** | **58** | **57** | **58** |
| **Strand (%)** | **3** | **7** | **3** | **8** | **5** | **8** |

**Supplementary Table S6**: Comparison of secondary structure content from SRCD, NMR and X-ray. NMR structure of apo-CaM is a consensus of 25 NMR structures (pdb 1CFC). NMR structure of Holo-CaM is a consensus of three structures of each lobe (pdb 1J7O). NMR and X-ray structures of Holo-CaM with CDZ correspond to the data presented in this paper (pdb 7PSZ, Holo-CaM:CDZ_A_ 1:1 and pdb 7PU9, Holo-CaM:CDZ_BC_ 1:2).
